## Supplemental File 1 for "Imputation of 3D genome structure by genetic-epigenetic interaction modeling in mice"

### STREME
Sensitive, Thorough, Rapid, Enriched Motif Elicitation

For further information on how to interpret these results please access https://meme-suite.org/meme/doc/streme.html.

To get a copy of the MEME software please access https://meme-suite.org.

If you use STREME in your research, please cite the following paper:

Timothy L. Bailey, "STREME: accurate and versatile sequence motif discovery", *Bioinformatics*, Mar. 24, 2021. [full text] If you use STREME in your research, please cite the following paper:
Timothy L. Bailey, "STREME: accurate and versatile sequence motif discovery", *Bioinformatics*, Mar. 24, 2021. [full text]

Discovered Motifs | Inputs & Settings | Program Information | Motifs in MEME Text Format | Matching Sequences NEW | Results in XML Format

#### Description

neg50 vs shuffled

#### Discovered Motifs

Next Top

| Motif | Logo | RC Logo | P-value | E-value NEW | Sites | More | Submit/Download | Positional Distribution NEW | Matches per Sequence NEW |
|---|---|---|---|---|---|---|---|---|---|
| 1-CCCCACCCCCACC | 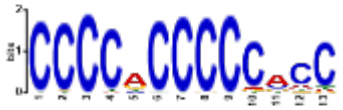 | 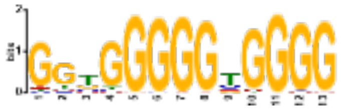 | 1.4e-022 | 7.1e-021 | 1338 (28.1%) | ↧ | ⇢ | 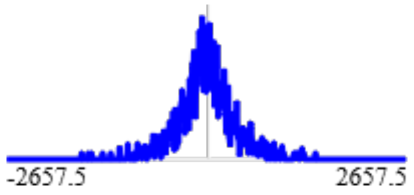  | 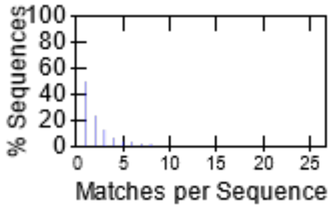  |
| 2-ACACACACACACACA | 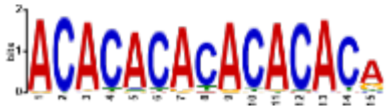 | 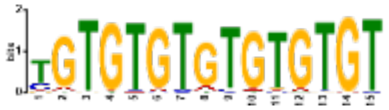 | 1.7e-014 | 8.6e-013 | 532 (11.2%) | ↧ | ⇢ | 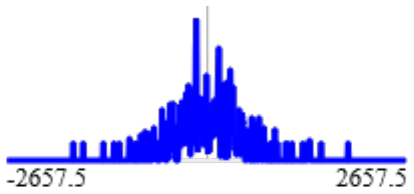  | 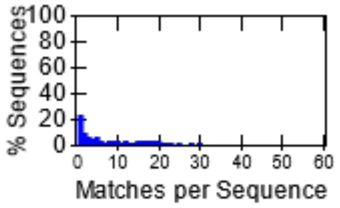  |
| 3-RCCASMAGAGGGCR | 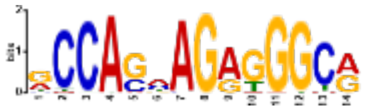 | 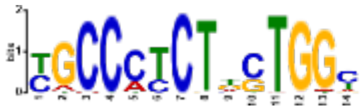 | 2.0e-014 | 1.0e-012 | 607 (12.8%) | ↧ | ⇢ | 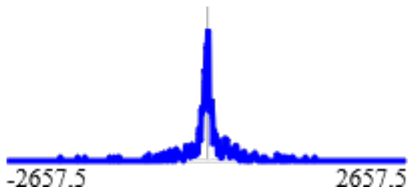  | 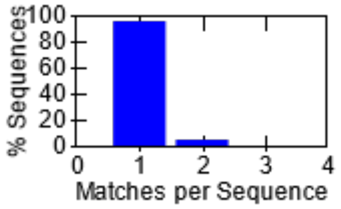  |
| 4-AAAAAAAAAAAAAAA | 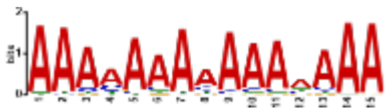 | 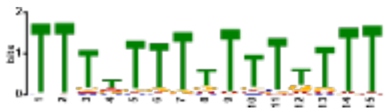 | 2.0e-014 | 1.0e-012 | 1102 (23.2%) | ↧ | ⇢ | 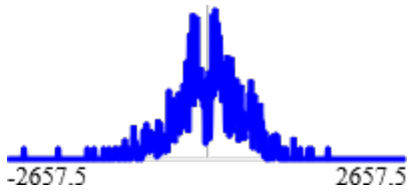  | 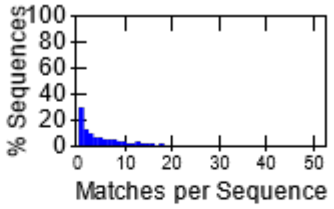  |

**Stopped because 3 consecutive motifs exceeded the p-value threshold (0.05).**
**STREME ran for 3043.51 seconds.**

| Motif | Logo | RC Logo | P-value | E-value NEW | Sites | More | Submit/Download | Positional Distribution NEW | Matches per Sequence NEW |
|---|---|---|---|---|---|---|---|---|---|
| 5-CCTGCCTCTGCCTC | 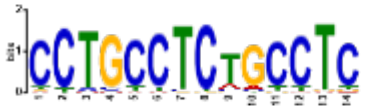 | 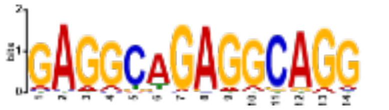 | 3.0e-012 | 1.5e-010 | 546 (11.5%) | | | 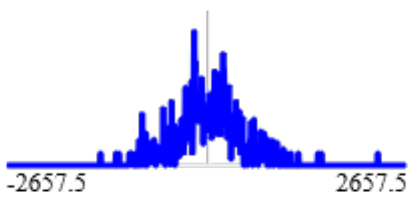 | 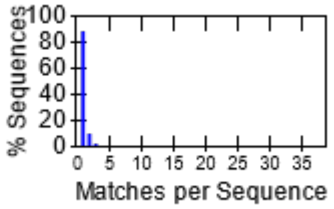 |
| 6-AGAGAGAGAGAGAGA | 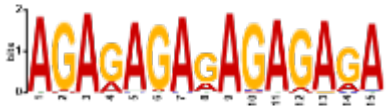 | 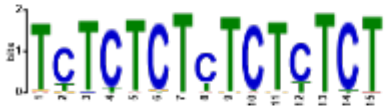 | 3.7e-012 | 1.9e-010 | 323 (6.8%) | | | 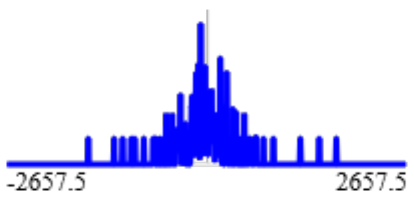 | 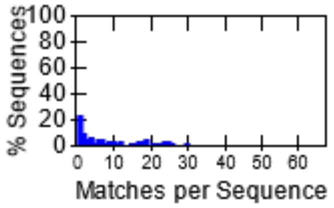 |
| 7-GAGTTCCAGGMCAGC | 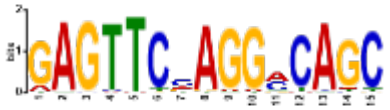 | 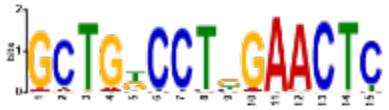 | 1.2e-010 | 6.3e-009 | 491 (10.3%) | | | 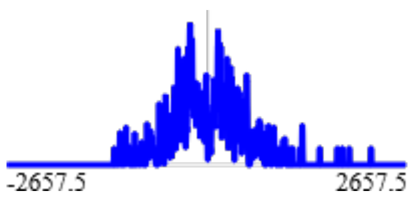 | 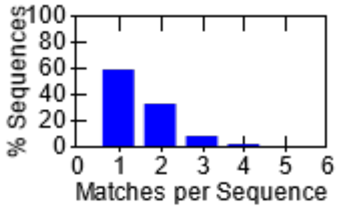 |
| 8-TGAGCCATCTCTCCA | 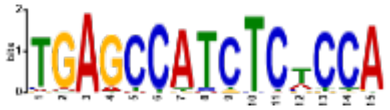 | 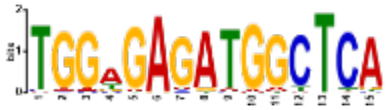 | 3.0e-008 | 1.6e-006 | 235 (4.9%) | | | 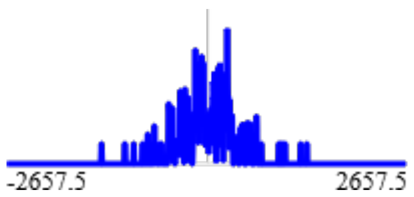 | 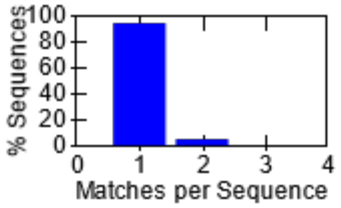 |
| 9-AGACAGGGTTTCTCT | 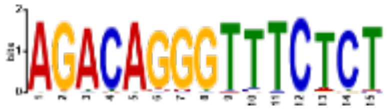 | 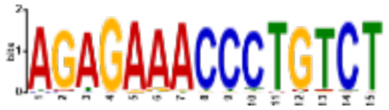 | 1.2e-007 | 6.2e-006 | 230 (4.8%) | | | 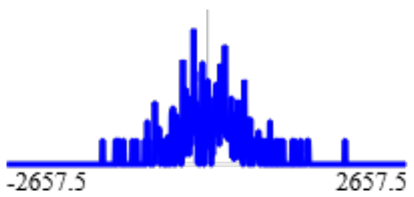 | 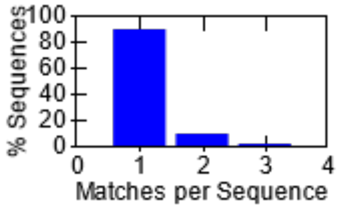 |
| 10-CCTTTAATCCCAGCA | 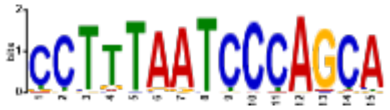 | 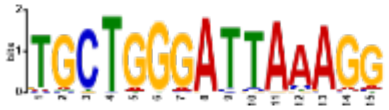 | 1.2e-007 | 6.4e-006 | 383 (8.0%) | | | 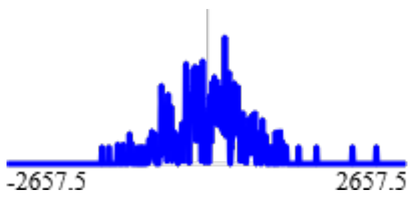 | 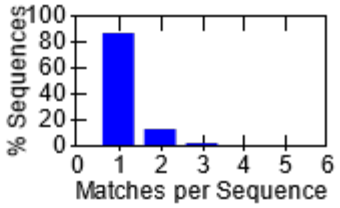 |
| 11-CCCACCCCACCCC | 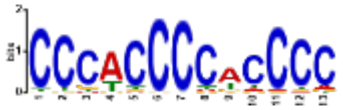 | 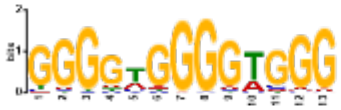 | 1.9e-007 | 1.0e-005 | 681 (14.3%) | | | 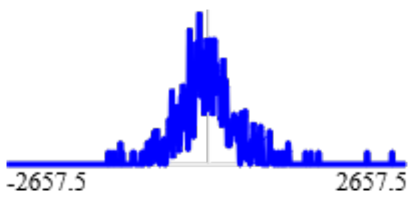 | 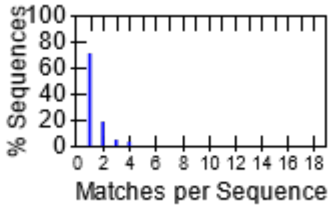 |

**Stopped because 3 consecutive motifs exceeded the p-value threshold (0.05).**
**STREME ran for 3043.51 seconds.**

| Motif | Logo | RC Logo | P-value | E-value NEW | Sites | More | Submit/Download | Positional Distribution NEW | Matches per Sequence NEW |
|---|---|---|---|---|---|---|---|---|---|
| 12-GGAGGGAGGGAGGGA | 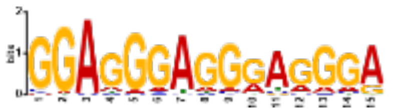 | 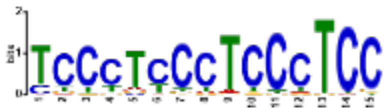 | 3.7e-007 | 1.9e-005 | 478 (10.0%) | ↧ | ⇢ | 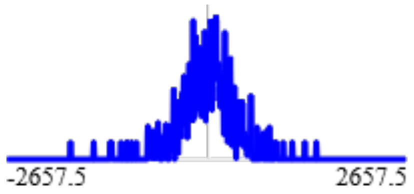 | 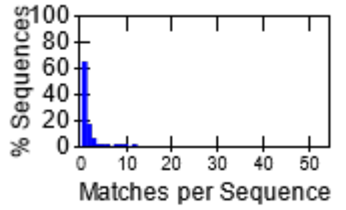 |
| 13-GCCCCGCCCC | 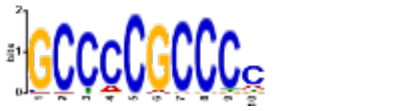 | 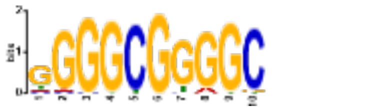 | 4.9e-007 | 2.5e-005 | 780 (16.4%) | ↧ | ⇢ | 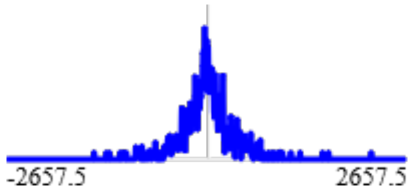 | 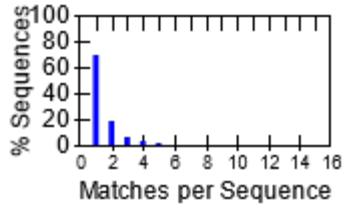 |
| 14-CRTGGTGGCKCACR | 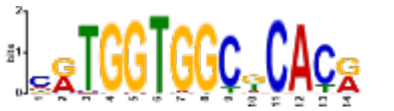 | 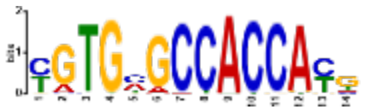 | 1.3e-006 | 6.7e-005 | 353 (7.4%) | ↧ | ⇢ | 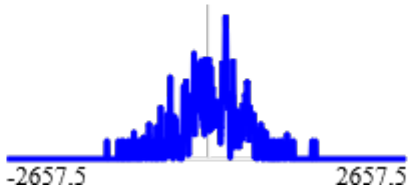 | 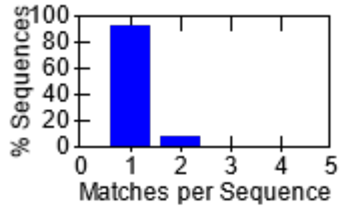 |
| 15-CTGGTCTACARAGT | 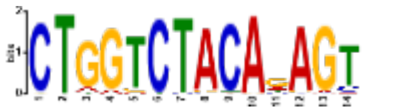 | 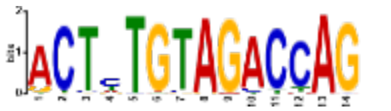 | 2.8e-006 | 1.5e-004 | 291 (6.1%) | ↧ | ⇢ | 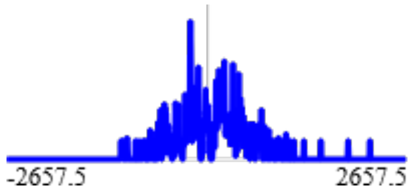 | 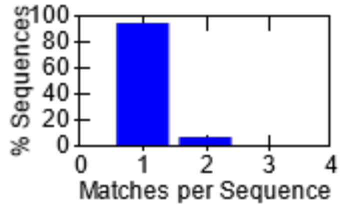 |
| 16-CGCCGCCGCCGCCG | 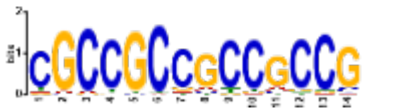 | 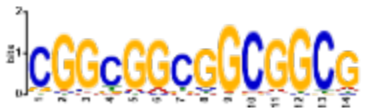 | 7.6e-006 | 4.0e-004 | 301 (6.3%) | ↧ | ⇢ | 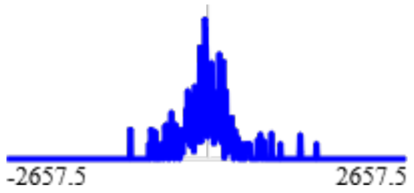 | 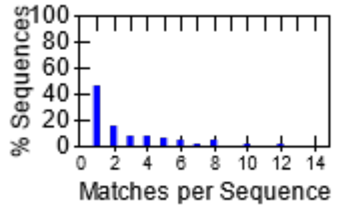 |
| 17-TGCTGGGAWTTGAAC | 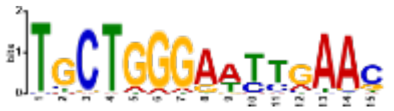 | 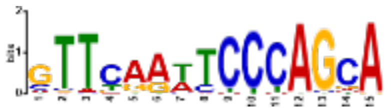 | 1.3e-005 | 6.6e-004 | 363 (7.6%) | ↧ | ⇢ | 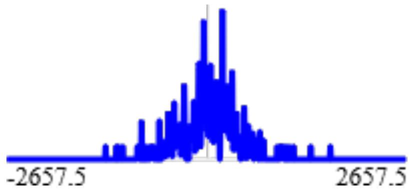 | 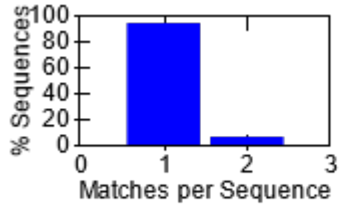 |
| 18-AGTGCTCTTAACCRC | 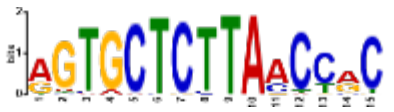 | 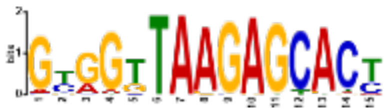 | 3.1e-005 | 1.6e-003 | 156 (3.3%) | ↧ | ⇢ | 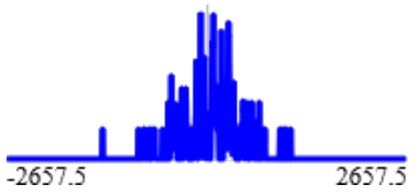 | 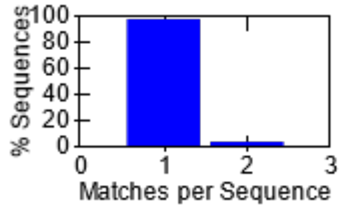 |

**Stopped because 3 consecutive motifs exceeded the p-value threshold (0.05).**
**STREME ran for 3043.51 seconds.**

| Motif | Logo | RC Logo | P-value | E-value NEW | Sites | More | Submit/Download | Positional Distribution NEW | Matches per Sequence NEW |
|---|---|---|---|---|---|---|---|---|---|
| 19-AAATAAATAT | 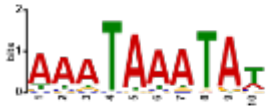 | 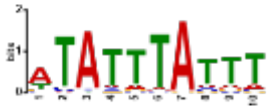 | 6.1e-005 | 3.2e-003 | 919 (19.3%) | ↧ | ⇢ | 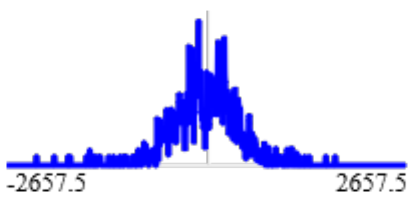  | 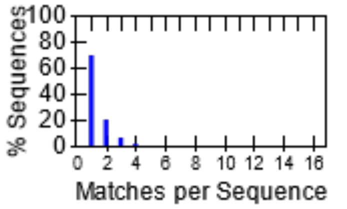  |
| 20-CAGGGCTAYAC | 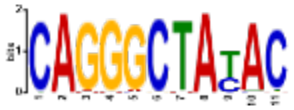 | 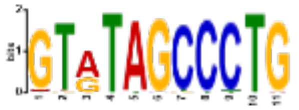 | 1.4e-004 | 7.2e-003 | 203 (4.3%) | ↧ | ⇢ | 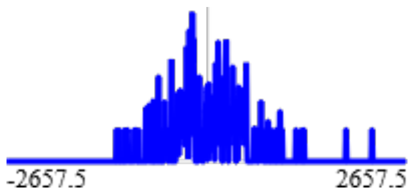  | 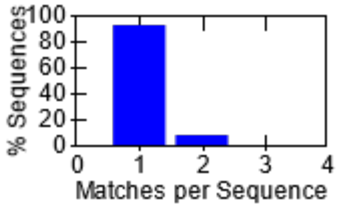  |
| 21-CCCCTCCCC | 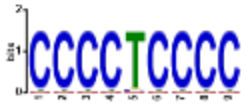 | 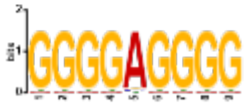 | 1.7e-004 | 8.8e-003 | 433 (9.1%) | ↧ | ⇢ | 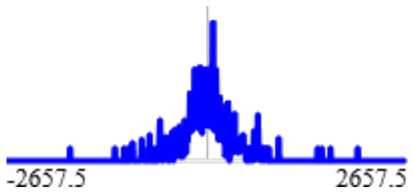  | 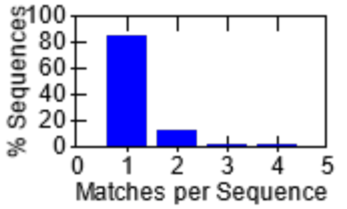  |
| 22-AGGAAGCAG | 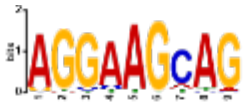 | 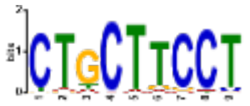 | 2.4e-004 | 1.3e-002 | 1332 (28.0%) | ↧ | ⇢ | 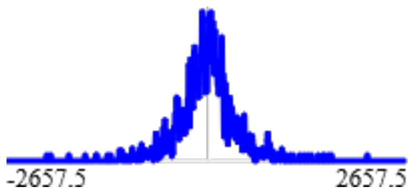  | 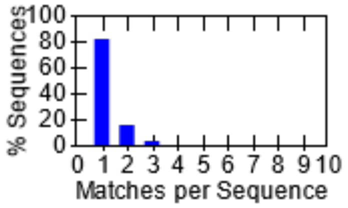  |
| 23-AAAACAAA | 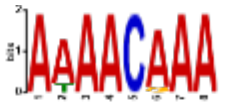 | 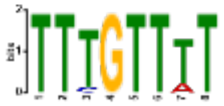 | 3.2e-004 | 1.6e-002 | 484 (10.2%) | ↧ | ⇢ | 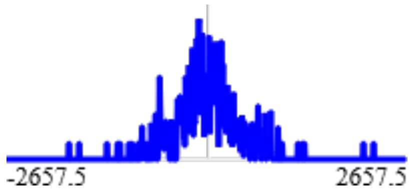  | 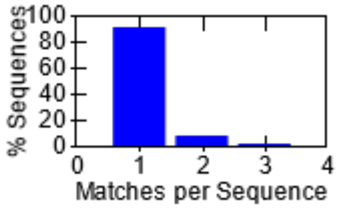  |
| 24-ATGCAAATD | 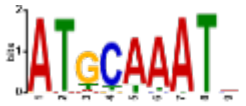 | 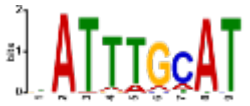 | 4.6e-004 | 2.4e-002 | 533 (11.2%) | ↧ | ⇢ | 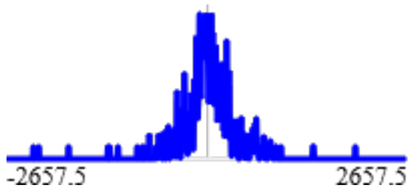  | 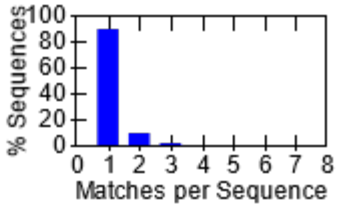  |
| 25-CCTYTGGAAGAGCAG | 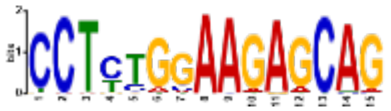 | 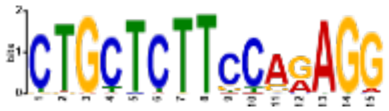 | 4.9e-004 | 2.5e-002 | 166 (3.5%) | ↧ | ⇢ | 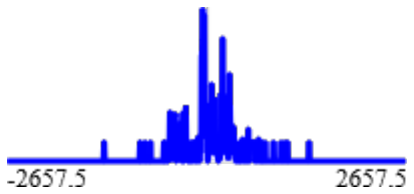  | 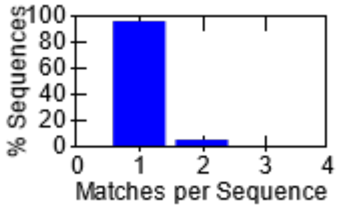  |

**Stopped because 3 consecutive motifs exceeded the p-value threshold (0.05).**
**STREME ran for 3043.51 seconds.**

| Motif | Logo | RC Logo | P-value | E-value NEW | Sites | More | Submit/Download | Positional Distribution NEW | Matches per Sequence NEW |
|---|---|---|---|---|---|---|---|---|---|
| 26-CCCTTCCCC | 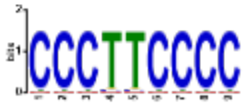 | 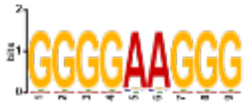 | 8.9e-004 | 4.6e-002 | 293 (6.2%) | ↧ | ⇢ | 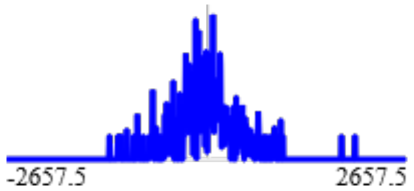 | 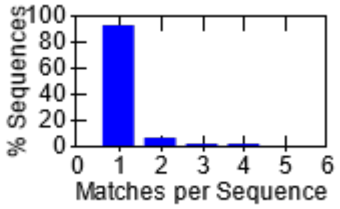 |
| 27-CTCCTCCTCCTCCYY | 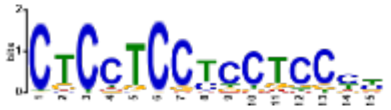 | 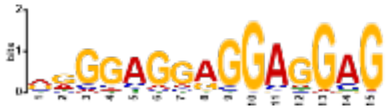 | 1.8e-003 | 9.2e-002 | 880 (18.5%) | ↧ | ⇢ | 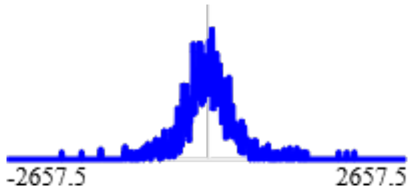 | 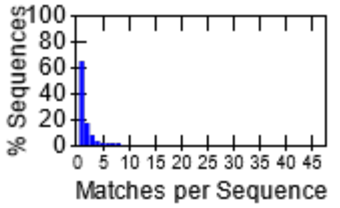 |
| 28-CTCCACCC | 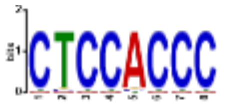 | 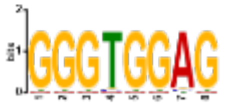 | 2.1e-003 | 1.1e-001 | 583 (12.3%) | ↧ | ⇢ | 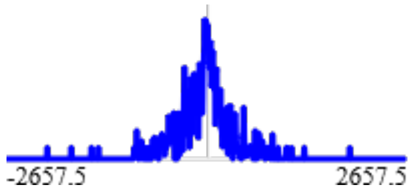 | 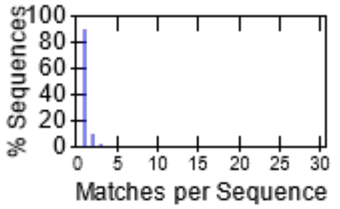 |
| 29-TTAAAAAWAAA | 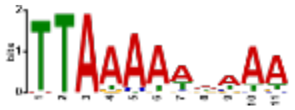 | 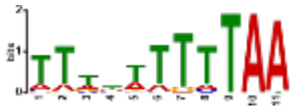 | 2.7e-003 | 1.4e-001 | 730 (15.3%) | ↧ | ⇢ | 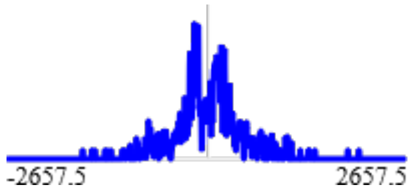 | 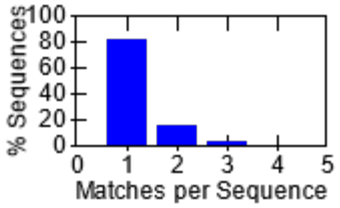 |
| 30-CGCGCGCGCGCG | 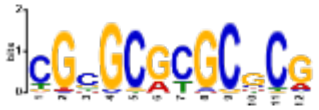 | 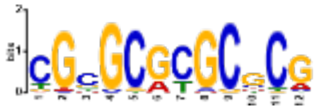 | 3.2e-003 | 1.7e-001 | 301 (6.3%) | ↧ | ⇢ | 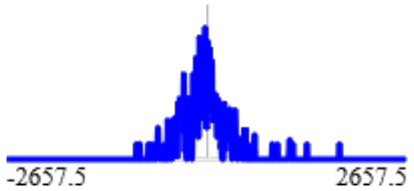 | 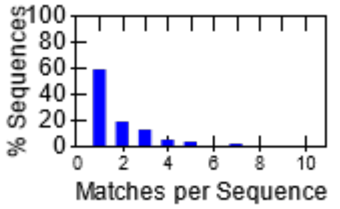 |
| 31-CCCGCCCCC | 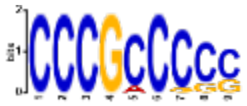 | 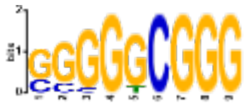 | 3.8e-003 | 2.0e-001 | 379 (8.0%) | ↧ | ⇢ | 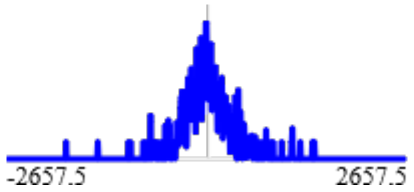 | 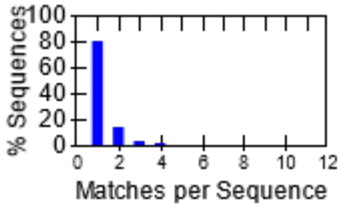 |
| 32-GTTACCTRGCAA | 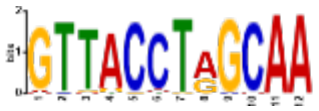 | 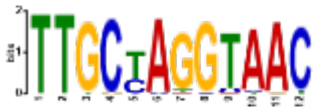 | 3.9e-003 | 2.0e-001 | 87 (1.8%) | ↧ | ⇢ | 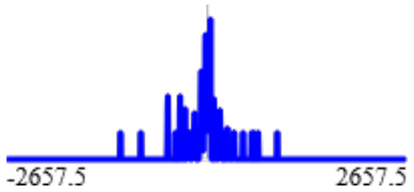 | 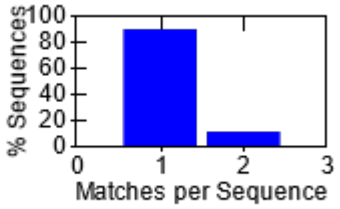 |

**Stopped because 3 consecutive motifs exceeded the p-value threshold (0.05).**
**STREME ran for 3043.51 seconds.**

| Motif | Logo | RC Logo | P-value | E-value | Sites | More | Submit/Download | Positional Distribution | Matches per Sequence |
|---|---|---|---|---|---|---|---|---|---|
| 33-ACCAATCAG | | | 5.8e-003 | 3.0e-001 | 272 (5.7%) | | | | 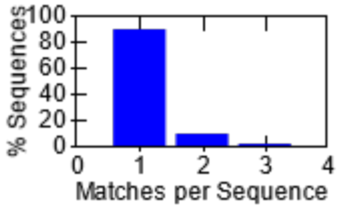 |
| 34-CGCCGCCGCC | | | 6.9e-003 | 3.6e-001 | 271 (5.7%) | | | | 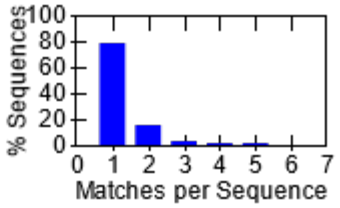 |
| 35-CAAGGTCAC | | | 1.0e-002 | 5.3e-001 | 505 (10.6%) | | | | 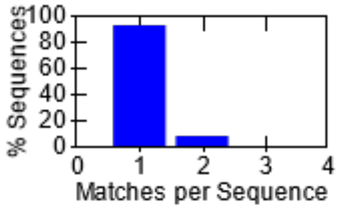 |
| 36-CAAAGCCA | | | 1.4e-002 | 7.5e-001 | 1848 (38.8%) | | | | 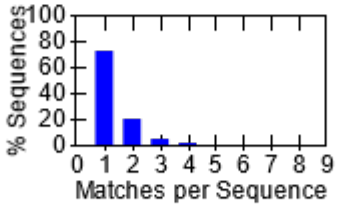 |
| 37-CCACCACCWCS | | | 1.5e-002 | 7.6e-001 | 335 (7.0%) | | | | 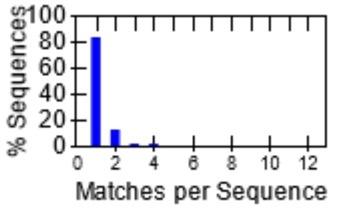 |
| 38-VTGASTCAB | | | 1.6e-002 | 8.2e-001 | 615 (12.9%) | | | | 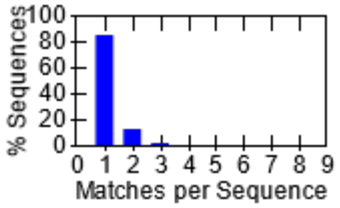 |
| 39-CAGCAGCTGC | | | 1.8e-002 | 9.2e-001 | 582 (12.2%) | | | | 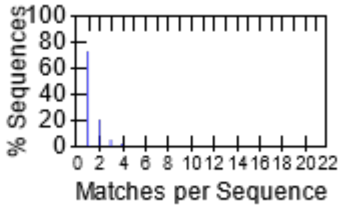 |

**Stopped because 3 consecutive motifs exceeded the p-value threshold (0.05).**
**STREME ran for 3043.51 seconds.**

| Motif | Logo | RC Logo | P-value | E-value | Sites | More | Submit/Download | Positional Distribution | Matches per Sequence |
|---|---|---|---|---|---|---|---|---|---|
| **40-AATGAATG** | | | 2.0e-002 | 1.0e+000 | 284 (6.0%) | ↧ | ⇢ | | 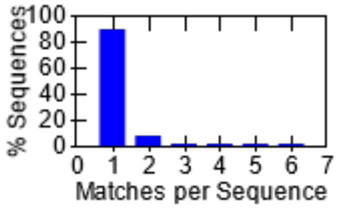 |
| **41-CAAGATGGCG** | | | 2.5e-002 | 1.3e+000 | 177 (3.7%) | ↧ | ⇢ | | 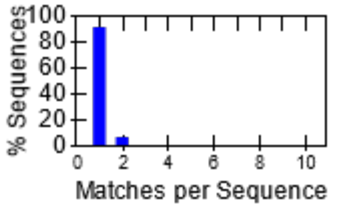 |
| **42-AAGAAAGAAA** | | | 3.4e-002 | 1.8e+000 | 456 (9.6%) | ↧ | ⇢ | | 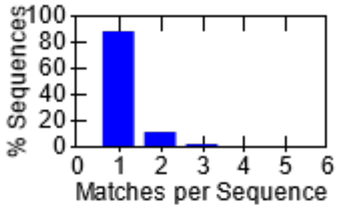 |
| **43-CCCAGCCC** | | | 3.6e-002 | 1.8e+000 | 752 (15.8%) | ↧ | ⇢ | | 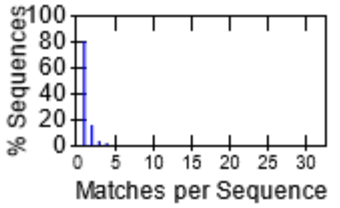 |
| **44-YCACGTGR** | | | 3.6e-002 | 1.9e+000 | 651 (13.7%) | ↧ | ⇢ | | 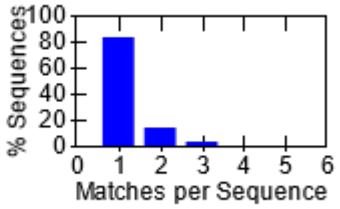 |
| **45-CCCCACCC** | | | 4.3e-002 | 2.2e+000 | 580 (12.2%) | ↧ | ⇢ | | 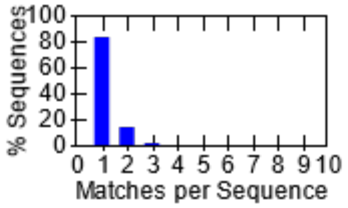 |
| **46-CATCTGTAA** | | | 4.5e-002 | 2.3e+000 | 179 (3.8%) | ↧ | ⇢ | | 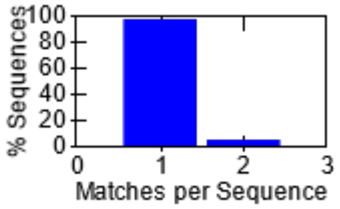 |

**Stopped because 3 consecutive motifs exceeded the p-value threshold (0.05).**
**STREME ran for 3043.51 seconds.**

| Motif | Logo | RC Logo | P-value | E-value NEW | Sites | More | Submit/Download | Positional Distribution NEW | Matches per Sequence NEW |
|---|---|---|---|---|---|---|---|---|---|
| 47-CCGCCTCC | | | 4.5e-002 | 2.4e+000 | 917 (19.3%) | ↧ | ⋯→ | | 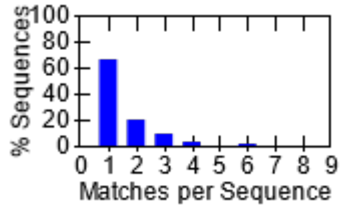 |
| 48-GACAGACA | | | 4.6e-002 | 2.4e+000 | 346 (7.3%) | ↧ | ⋯→ | | 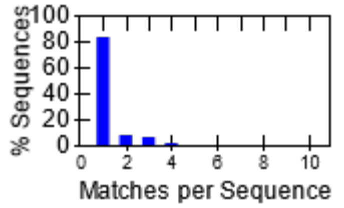 |
| 49-GCGCAGGCGCA | | | 4.8e-002 | 2.5e+000 | 146 (3.1%) | ↧ | ⋯→ | | 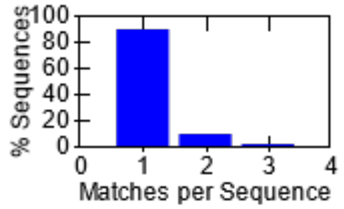 |
| 50-AACAACAACAA | | | 9.0e-002 | 4.7e+000 | 117 (2.5%) | ↧ | ⋯→ | | 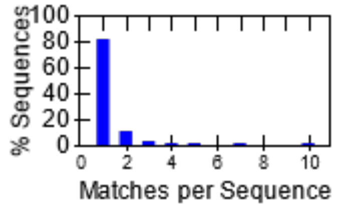 |
| 51-AGCCTGGGCTACA | | | 1.1e-001 | 5.9e+000 | 172 (3.6%) | ↧ | ⋯→ | | 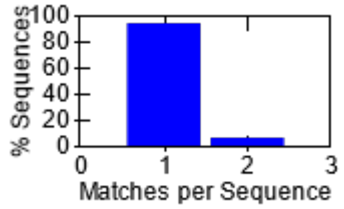 |
| 52-ACTGTAGCTGTCTTC | | | 1.9e-001 | 9.8e+000 | 85 (1.8%) | ↧ | ⋯→ | | 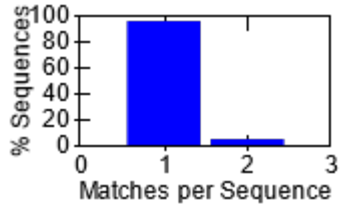 |

**Stopped because 3 consecutive motifs exceeded the p-value threshold (0.05).**
**STREME ran for 3043.51 seconds.**

#### Inputs & Settings

Previous Next Top

##### Sequences

| Role | Source | Alphabet | Sequence Count | Total Size |
|---|---|---|---|---|
| Positive (primary) Sequences | neg50_atac_cleaned_20211005.fasta | DNA | 4758 | 3999975 |
| Negative (control) Sequences | 2-Order Shuffled Positive Sequences | DNA | 4778 | 3999890 |

##### Background Model

**Source:** built from the negative (control) sequences

**Source:** built from the negative (control) sequences

**Order:** 2 (only order-0 shown)

| Name | Freq. | Bg. | | | | Bg. | Freq. | Name |
|---|---|---|---|---|---|---|---|---|
| Adenine | 0.229 | 0.229 | A | ~ | T | 0.229 | 0.229 | Thymine |
| Cytosine | 0.271 | 0.271 | C | ~ | G | 0.271 | 0.271 | Guanine |


| Name | Freq. | Bg. | | | | Bg. | Freq. | Name |
|---|---|---|---|---|---|---|---|---|
| Adenine | 0.229 | 0.229 | A | ~ | T | 0.229 | 0.229 | Thymine |
| Cytosine | 0.271 | 0.271 | C | ~ | G | 0.271 | 0.271 | Guanine |


#### Other Settings

| | |
|---|---|
| **Strand Handling** | Both the given and reverse complement strands are processed. |
| **Objective Function** | Differential Enrichment |
| **Statistical Test** | Binomial Test |
| **Minimum Motif Width** | 8 |
| **Maximum Motif Width** | 15 |
| **Sequence Shuffling** | Negative sequences are positives shuffled preserving 3-mer frequencies. |
| **Test Set** | 10% of the input sequences were randomly assigned to the test set. |
| **Word Evaluation** | Up to 25 words of each width from 8 to 15 were evaluated to find seeds. |
| **Seed Refinement** | Up to 4 seeds of each width from 8 to 15 were further refined. |
| **Refinement Iterations** | Up to 20 iterations were allowed when refining a seed. |
| **Random Number Seed** | 0 |
| **Total Length** | The total length of each sequence set was limited to 4.00e+6. |
| **Maximum Motif p-value** | Stop when the p-value is greater than 0.05 for 3 consecutive motifs. |
| **Maximum Motifs to Find** | No maximum number of motifs. |
| **Maximum Run Time** | 14400 seconds. |

Previous Top

**STREME version**
5.4.1 (Release date: Sat Aug 21 19:23:23 2021 -0700)

**Reference**
Timothy L. Bailey, "STREME: accurate and versatile sequence motif discovery", *Bioinformatics*, Mar. 24, 2021. [full text]

**Reference**
Timothy L. Bailey, "STREME: accurate and versatile sequence motif discovery", *Bioinformatics*, Mar. 24, 2021. [full text]

**Command line**

```
streme --verbosity 1 --oc . --dna --totallength 4000000 --time 14400 --minw 8 --maxw 15 --thresh 0.05 --align center --dfile description --p neg50_atac_cleaned_20211005.fasta
```
