## Supplemental File 2 for "Imputation of 3D genome structure by genetic-epigenetic interaction modeling in mice"

### Tomtom
Motif Comparison Tool

For further information on how to interpret these results please access https://meme-suite.org/meme/doc/tomtom-output-format.html.
To get a copy of the MEME software please access https://meme-suite.org.

If you use Tomtom in your research, please cite the following paper:
Shobhit Gupta, JA Stamatoyannopolous, Timothy Bailey and William Stafford Noble, "Quantifying similarity between motifs", *Genome Biology*, **8**(2):R24, 2007. [full text]

Query Motifs | Target Databases | Matches | Settings | Program information | Results in TSV Format | Results in XML Format

#### Query Motifs

Next Top

| Database | ID | Alt. ID | Preview | Matches | List |
|---|---|---|---|---|---|
| query_motifs | GTAAGACTCACTGGCCAATCAAGGTCKCCAGGCAGACATGCCAG | MEME-4 | 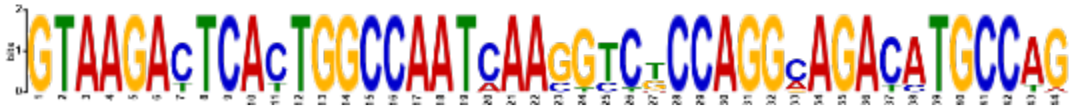 | 17 | SMAD2_MOUSE.H11MO.0.A, FOXI1_MOUSE.H11MO.0.B, |

#### Target Databases

Previous Next Top

| Database | Used | Matched |
|---|---|---|
| HOCOMOCOv11_core_MOUSE_mono_meme_format | 358 | 17 |

#### Matches to GTAAGACTCACTGGCCAATCAAGGTCKCCAGGCAGACATGCCAG (MEME-4)

Previous Next Top

**Summary**

| | |
|---|---|
| **Name** | SMAD2_MOUSE.H11MO.0.A |
| **Database** | HOCOMOCOv11_core_MOUSE_mono_meme_format |
| ***p*-value** | 2.43e-04 |
| ***E*-value** | 8.72e-02 |
| ***q*-value** | 1.74e-01 |
| **Overlap** | 9 |
| **Offset** | -27 |
| **Orientation** | Reverse Complement |
| | Show logo download options |

**Optimal Alignment**

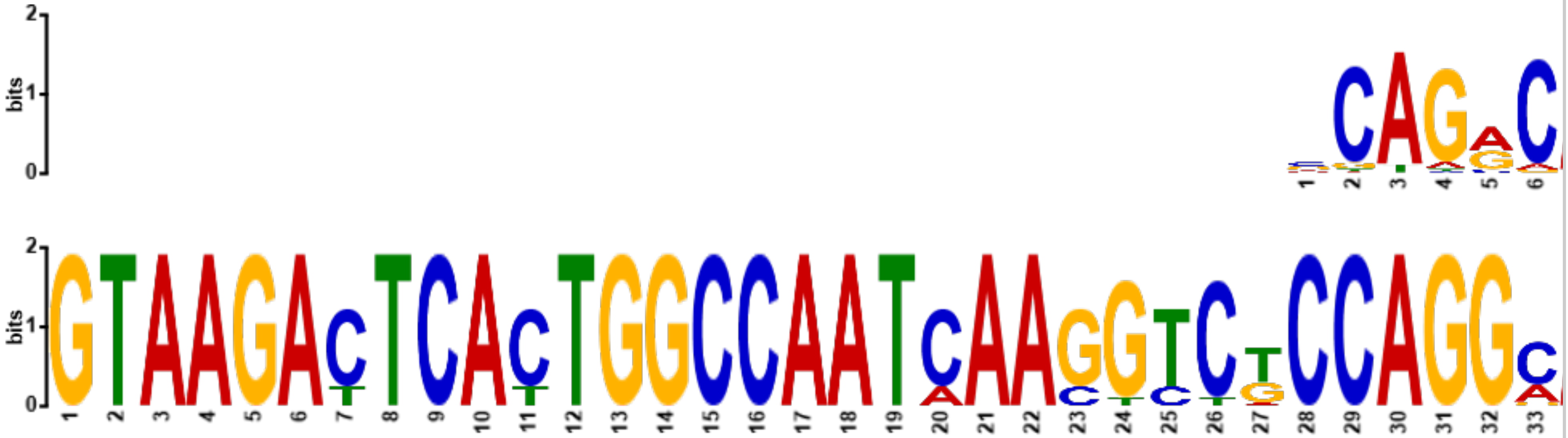


**Summary**

**Optimal Alignment**

| | |
|---|---|
| **Name** | FOXI1_MOUSE.H11MO.0.B |
| **Database** | HOCOMOCOv11_core_MOUSE_mono_meme_format |
| ***p*-value** | 1.54e-03 |
| ***E*-value** | 5.51e-01 |
| ***q*-value** | 3.92e-01 |
| **Overlap** | 12 |
| **Offset** | -12 |
| **Orientation** | Reverse Complement |

Show logo download options

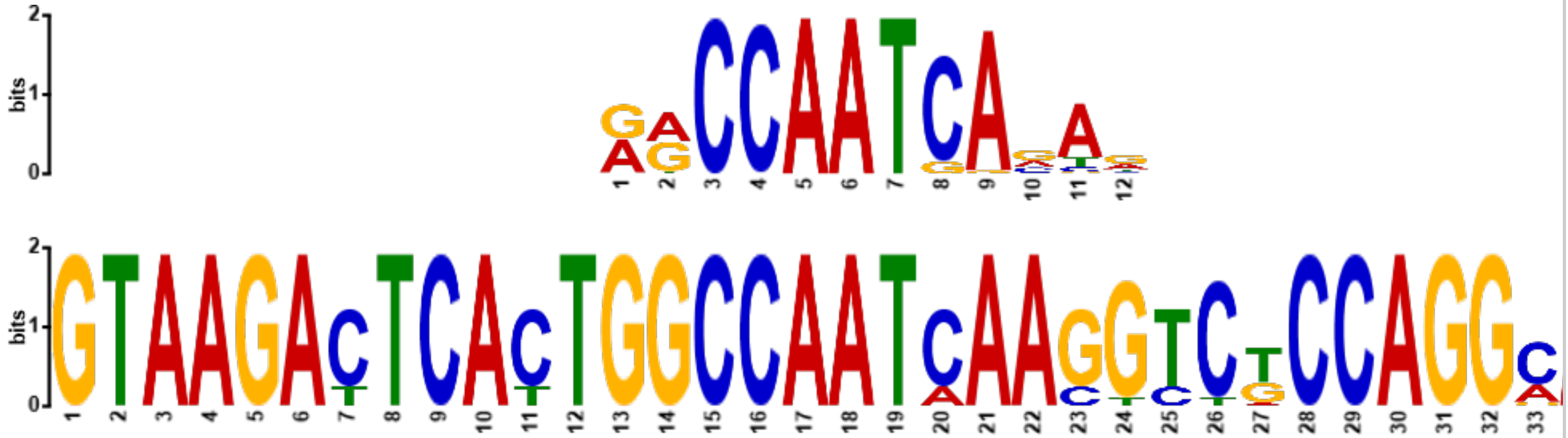

**Summary**

**Optimal Alignment**

| | |
|---|---|
| **Name** | NFYA_MOUSE.H11MO.0.A |
| **Database** | HOCOMOCOv11_core_MOUSE_mono_meme_format |
| ***p*-value** | 2.33e-03 |
| ***E*-value** | 8.35e-01 |
| ***q*-value** | 3.92e-01 |
| **Overlap** | 14 |
| **Offset** | -10 |
| **Orientation** | Normal |

Show logo download options

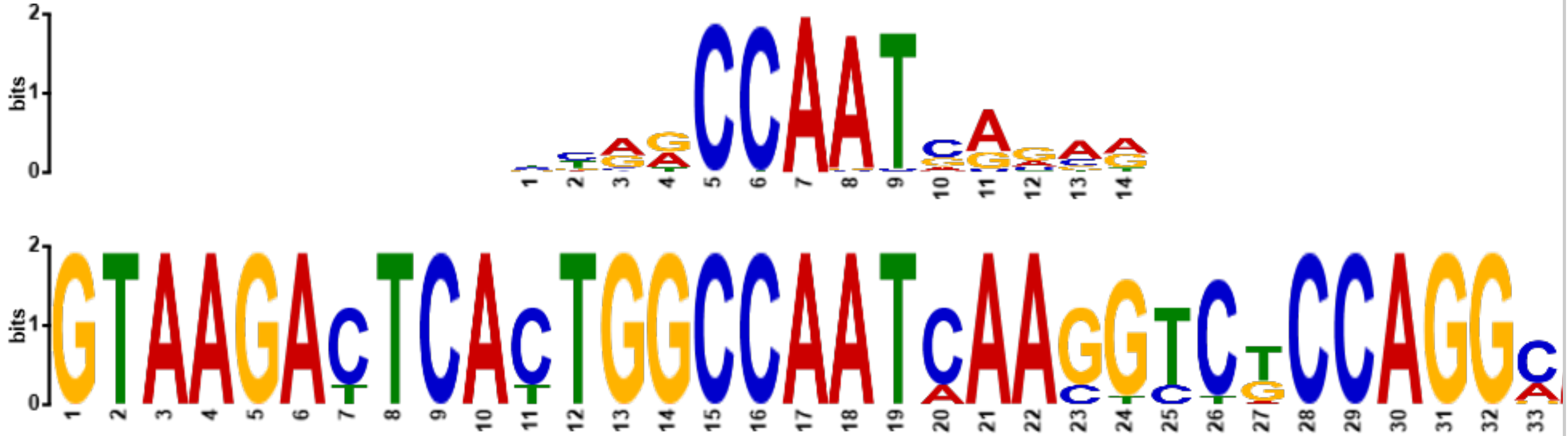

**Summary**

**Optimal Alignment**

| | |
|---|---|
| **Name** | ERR3_MOUSE.H11MO.0.C |
| **Database** | HOCOMOCOv11_core_MOUSE_mono_meme_format |
| ***p*-value** | 2.44e-03 |
| ***E*-value** | 8.75e-01 |
| ***q*-value** | 3.92e-01 |
| **Overlap** | 10 |
| **Offset** | -18 |
| **Orientation** | Normal |

Show logo download options

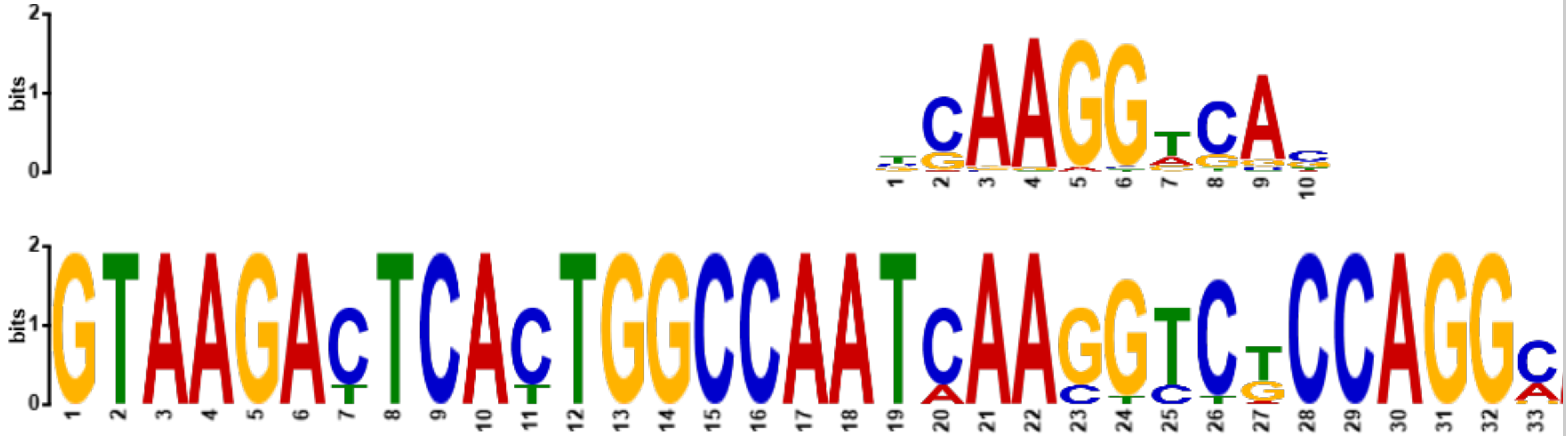

**Summary**

**Optimal Alignment**

**Name** [NFYC_MOUSE.H11MO.0.B]
**Database** HOCOMOCOv11_core_MOUSE_mono_meme_format

***p*-value** 2.90e-03
***E*-value** 1.04e+00
***q*-value** 3.92e-01

**Overlap** 14
**Offset** -10
**Orientation** Normal

Show logo download options

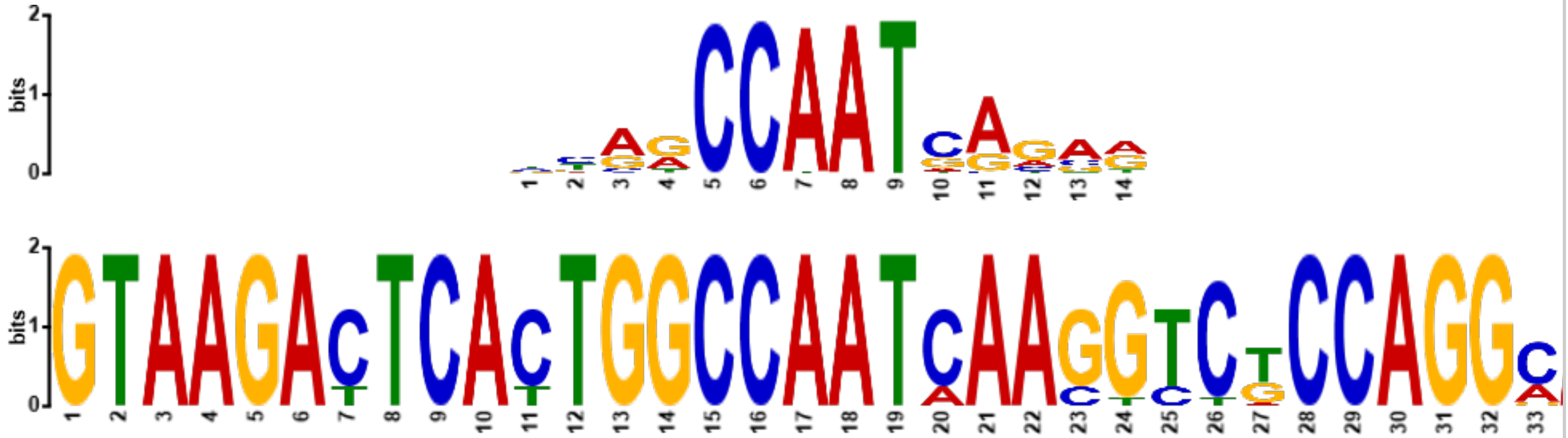

##### Summary

##### Optimal Alignment

**Name** [PBX3_MOUSE.H11MO.0.A]
**Database** HOCOMOCOv11_core_MOUSE_mono_meme_format

***p*-value** 3.30e-03
***E*-value** 1.18e+00
***q*-value** 3.92e-01

**Overlap** 14
**Offset** -10
**Orientation** Reverse Complement

Show logo download options

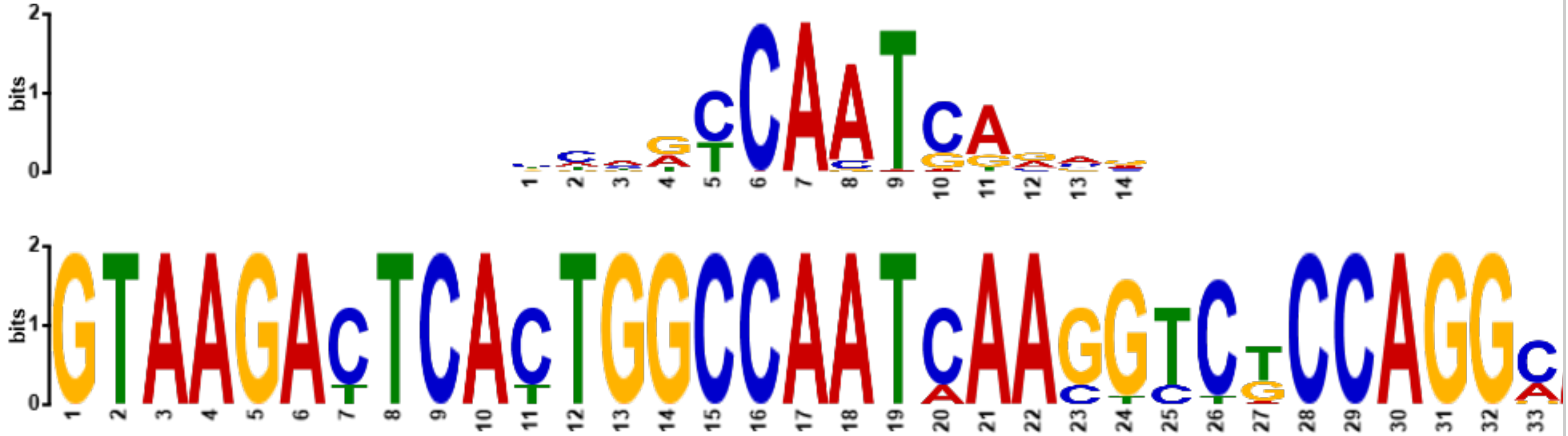

##### Summary

##### Optimal Alignment

**Name** [NFYB_MOUSE.H11MO.0.A]
**Database** HOCOMOCOv11_core_MOUSE_mono_meme_format

***p*-value** 5.60e-03
***E*-value** 2.01e+00
***q*-value** 5.71e-01

**Overlap** 13
**Offset** -11
**Orientation** Normal

Show logo download options

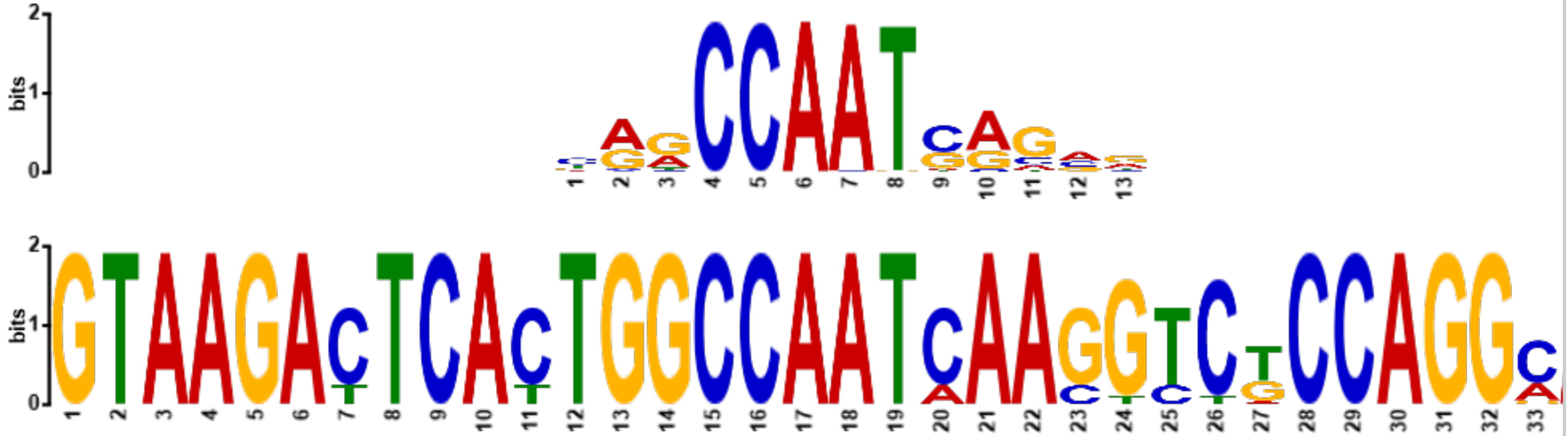

##### Summary

##### Optimal Alignment

| | |
|---|---|
| **Name** | CTCF_MOUSE.H11MO.0.A |
| **Database** | HOCOMOCOv11_core_MOUSE_mono_meme_format |
| ***p*-value** | 1.33e-02 |
| ***E*-value** | 4.76e+00 |
| ***q*-value** | 9.97e-01 |
| **Overlap** | 20 |
| **Offset** | -11 |
| **Orientation** | Normal |
| | Show logo download options |

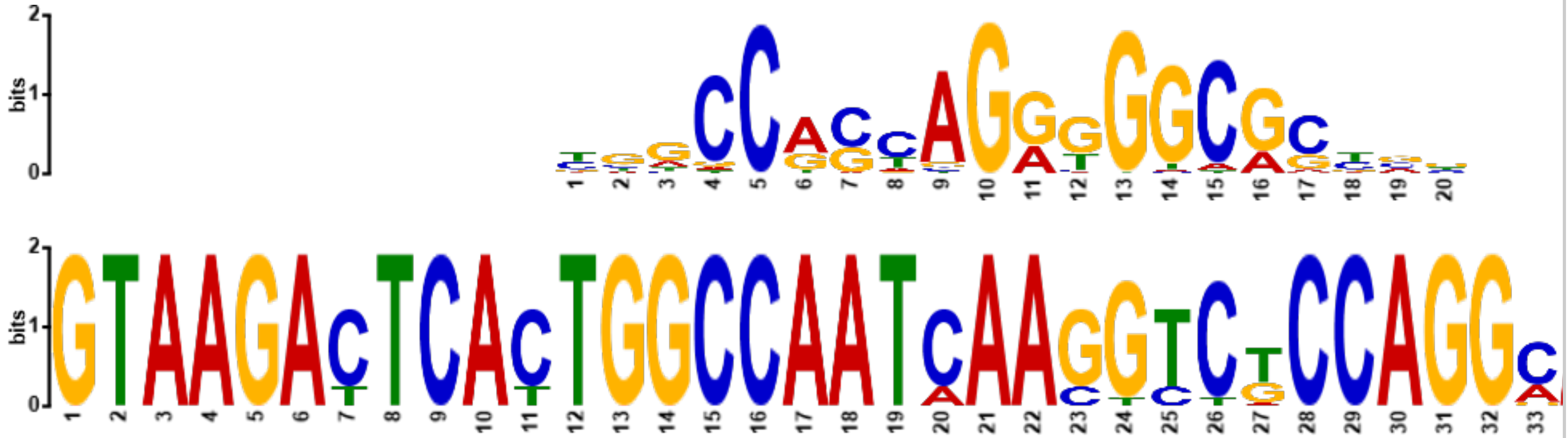

**Summary** **Optimal Alignment**

| | |
|---|---|
| **Name** | FOSB_MOUSE.H11MO.0.A |
| **Database** | HOCOMOCOv11_core_MOUSE_mono_meme_format |
| ***p*-value** | 1.39e-02 |
| ***E*-value** | 4.97e+00 |
| ***q*-value** | 9.97e-01 |
| **Overlap** | 9 |
| **Offset** | -2 |
| **Orientation** | Reverse Complement |
| | Show logo download options |

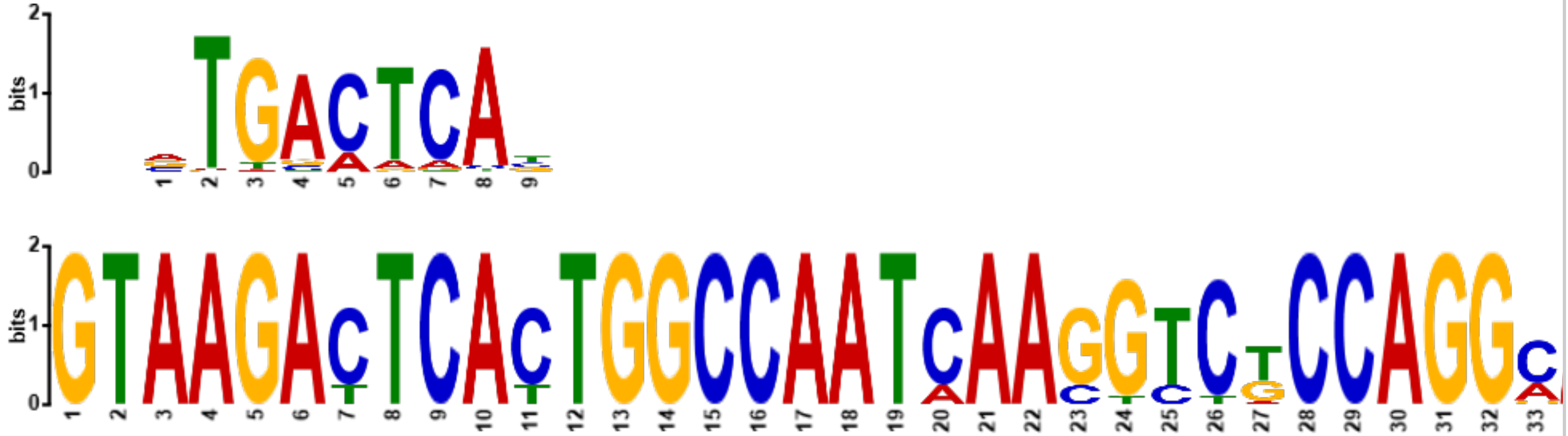

**Summary** **Optimal Alignment**

| | |
|---|---|
| **Name** | NKX21_MOUSE.H11MO.0.A |
| **Database** | HOCOMOCOv11_core_MOUSE_mono_meme_format |
| ***p*-value** | 1.62e-02 |
| ***E*-value** | 5.80e+00 |
| ***q*-value** | 9.97e-01 |
| **Overlap** | 10 |
| **Offset** | -14 |
| **Orientation** | Reverse Complement |
| | Show logo download options |

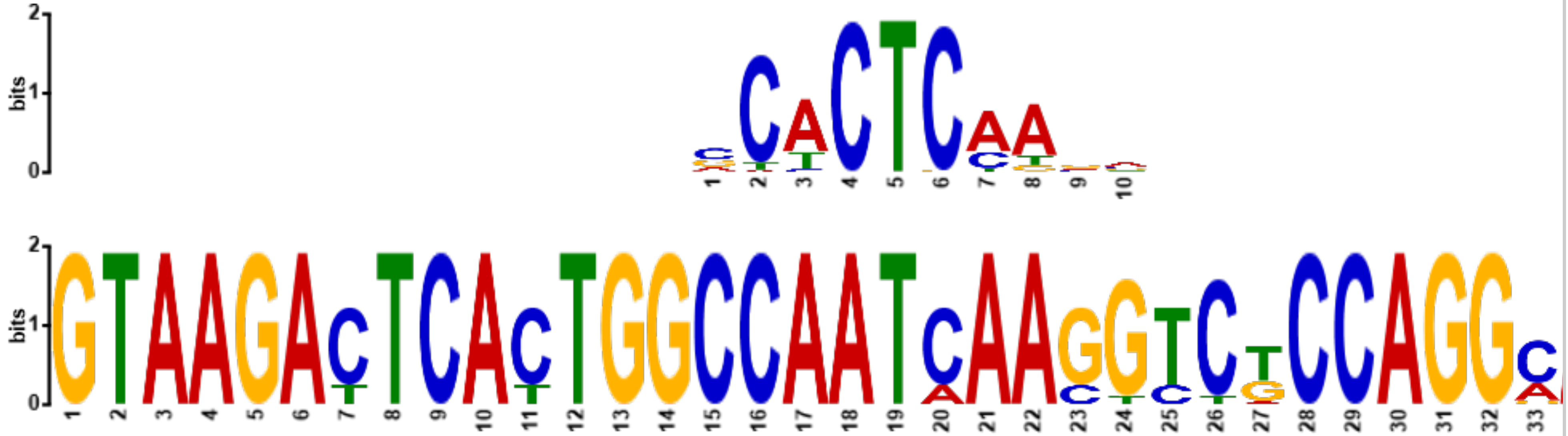

**Summary** **Optimal Alignment**

| | |
|---|---|
| **Name** | JUN_MOUSE.H11MO.0.A |
| **Database** | HOCOMOCOv11_core_MOUSE_mono_meme_format |
| ***p*-value** | 1.70e-02 |
| ***E*-value** | 6.08e+00 |
| ***q*-value** | 9.97e-01 |
| **Overlap** | 10 |
| **Offset** | -2 |
| **Orientation** | Reverse Complement |

Show logo download options

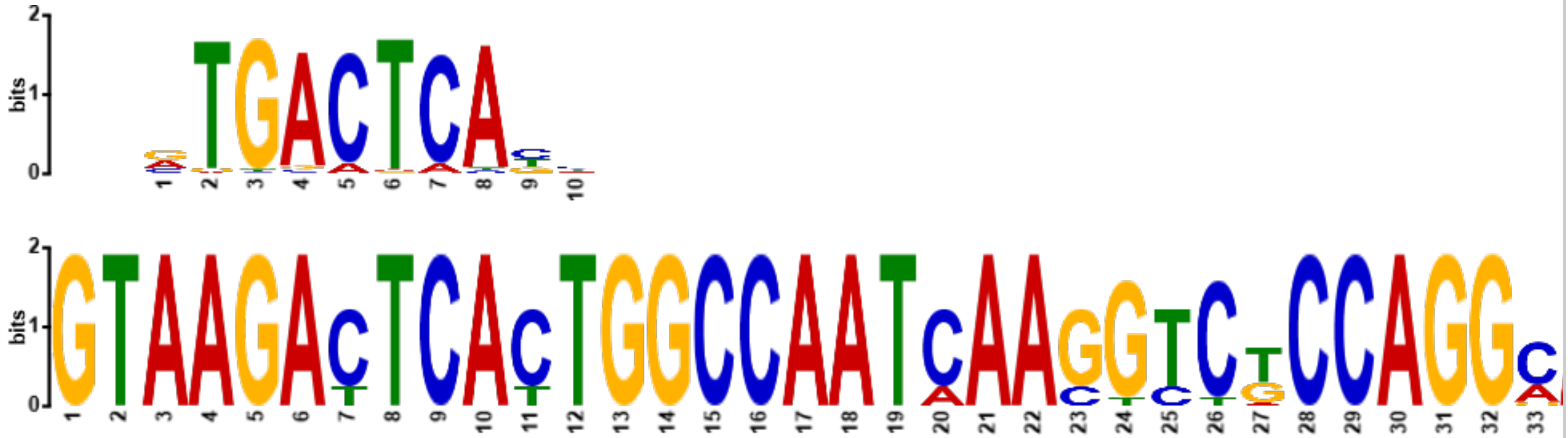

**Summary**

**Optimal Alignment**

| | |
|---|---|
| **Name** | JUNB_MOUSE.H11MO.0.A |
| **Database** | HOCOMOCOv11_core_MOUSE_mono_meme_format |
| ***p*-value** | 1.96e-02 |
| ***E*-value** | 7.01e+00 |
| ***q*-value** | 9.97e-01 |
| **Overlap** | 9 |
| **Offset** | -2 |
| **Orientation** | Reverse Complement |

Show logo download options

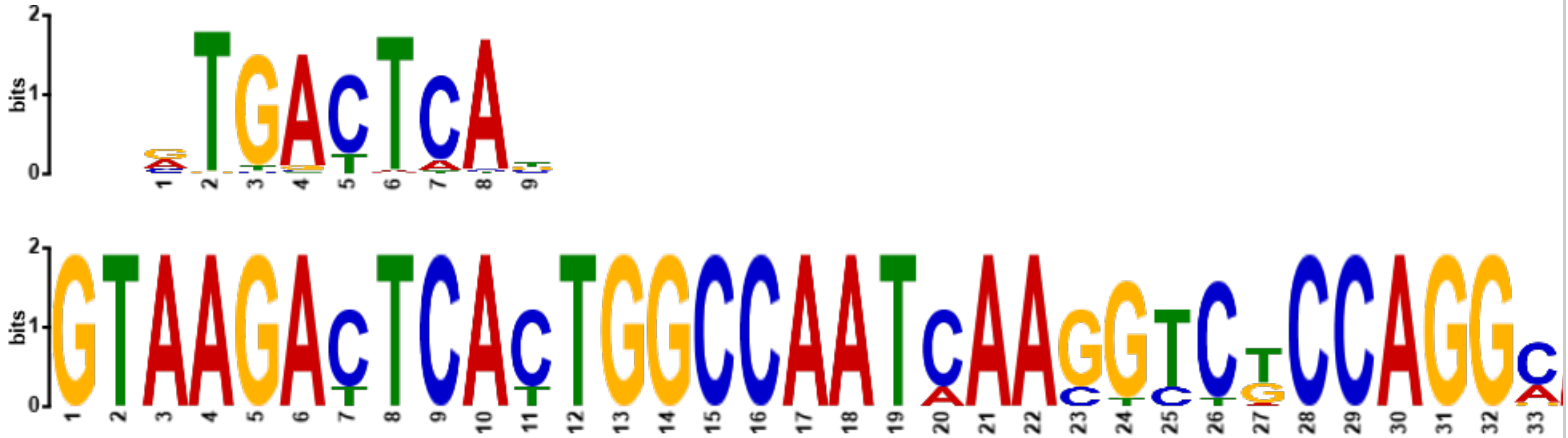

**Summary**

**Optimal Alignment**

| | |
|---|---|
| **Name** | JUND_MOUSE.H11MO.0.A |
| **Database** | HOCOMOCOv11_core_MOUSE_mono_meme_format |
| ***p*-value** | 2.16e-02 |
| ***E*-value** | 7.74e+00 |
| ***q*-value** | 9.97e-01 |
| **Overlap** | 11 |
| **Offset** | -2 |
| **Orientation** | Reverse Complement |

Show logo download options

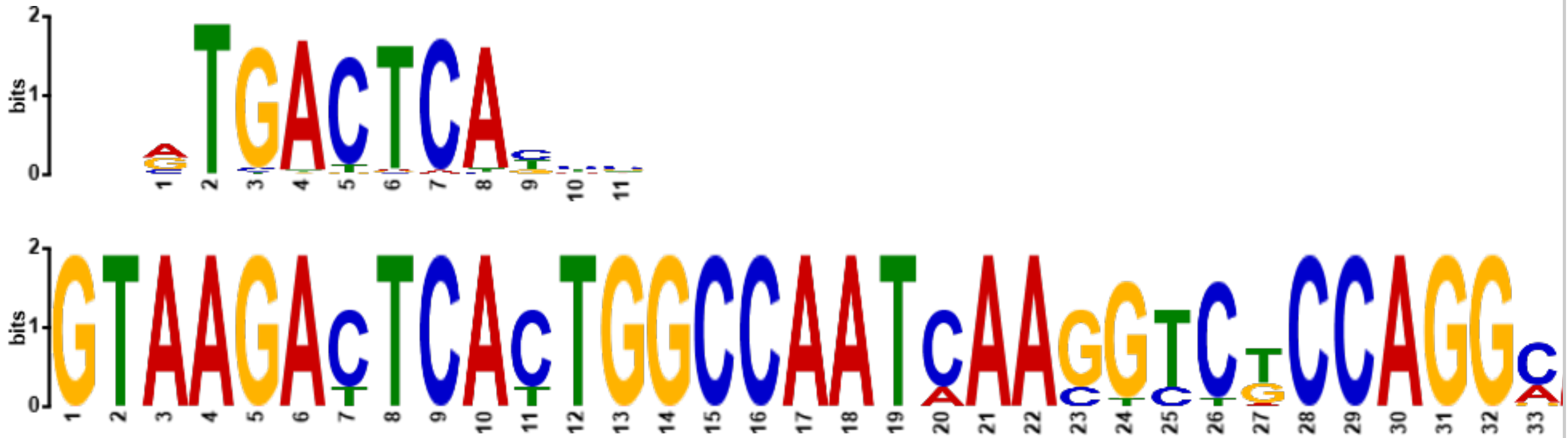

**Summary**

**Optimal Alignment**

| | |
|---|---|
| **Name** | BCL6_MOUSE.H11MO.0.A |
| **Database** | HOCOMOCOv11_core_MOUSE_mono_meme_format |
| ***p*-value** | 2.23e-02 |
| ***E*-value** | 7.98e+00 |
| ***q*-value** | 9.97e-01 |
| **Overlap** | 13 |
| **Offset** | -21 |
| **Orientation** | Normal |

Show logo download options

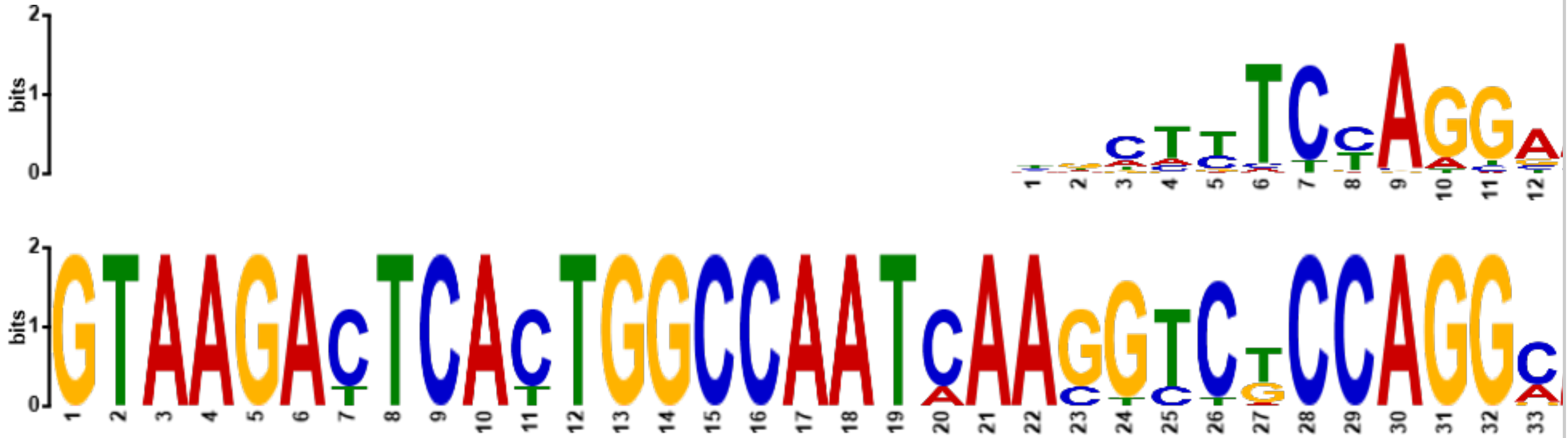

##### Summary

##### Optimal Alignment

| | |
|---|---|
| **Name** | ATF3_MOUSE.H11MO.0.A |
| **Database** | HOCOMOCOv11_core_MOUSE_mono_meme_format |
| ***p*-value** | 2.28e-02 |
| ***E*-value** | 8.15e+00 |
| ***q*-value** | 9.97e-01 |
| **Overlap** | 9 |
| **Offset** | -2 |
| **Orientation** | Reverse Complement |

Show logo download options

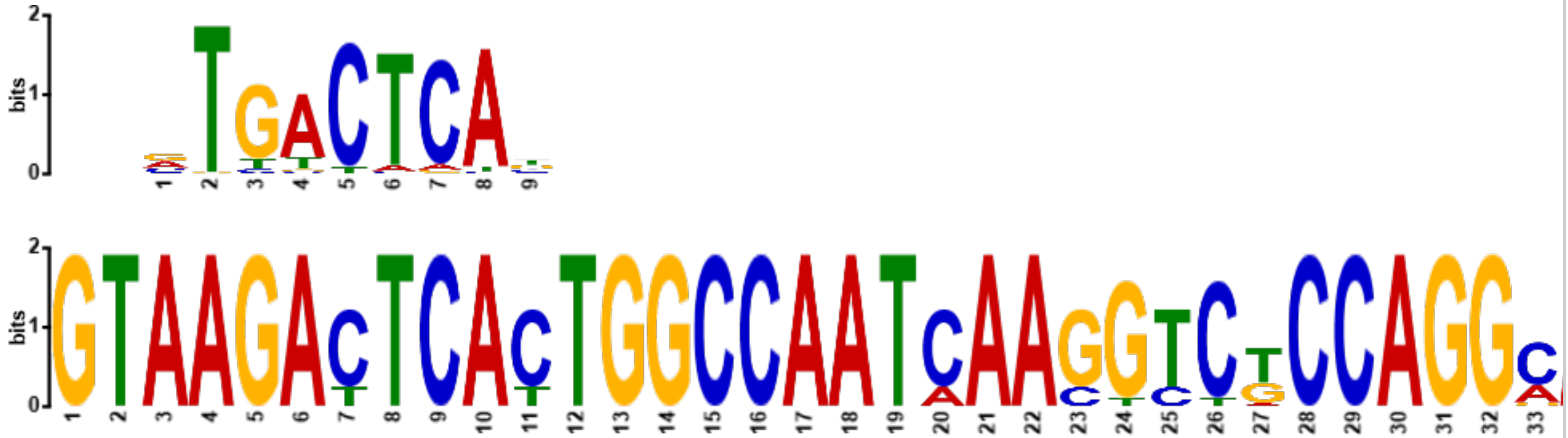

##### Summary

##### Optimal Alignment

| | |
|---|---|
| **Name** | TEAD1_MOUSE.H11MO.0.A |
| **Database** | HOCOMOCOv11_core_MOUSE_mono_meme_format |
| ***p*-value** | 2.68e-02 |
| ***E*-value** | 9.59e+00 |
| ***q*-value** | 9.97e-01 |
| **Overlap** | 9 |
| **Offset** | -35 |
| **Orientation** | Normal |

Show logo download options

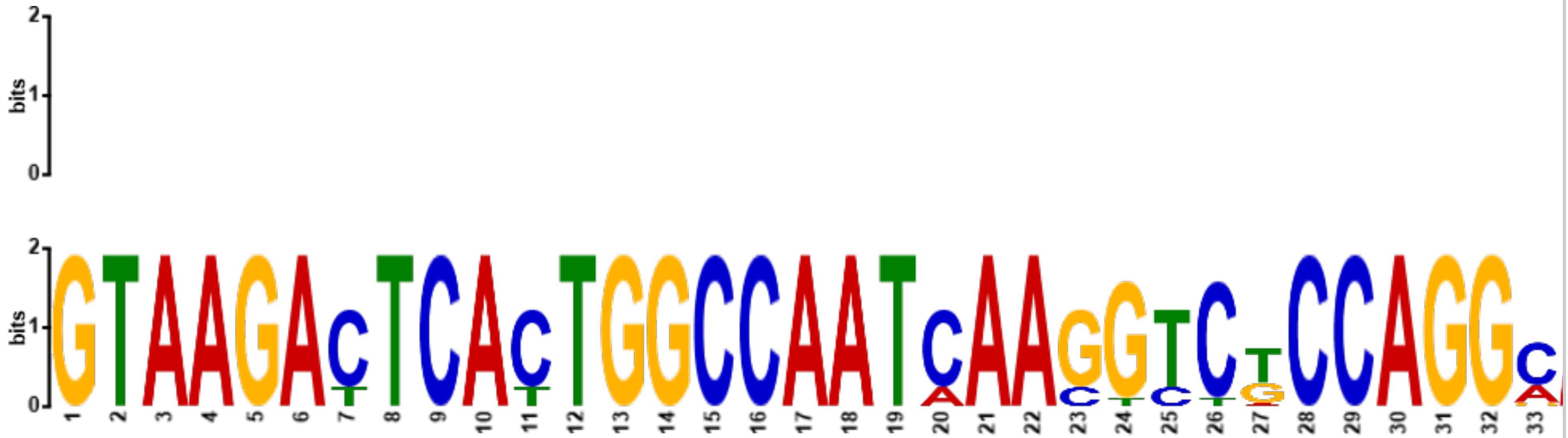

##### Summary

##### Optimal Alignment

| | |
|---|---|
| **Name** | LHX6_MOUSE.H11MO.0.C |
| **Database** | HOCOMOCOv11_core_MOUSE_mono_meme_format |
| ***p*-value** | 2.78e-02 |
| ***E*-value** | 9.96e+00 |
| ***q*-value** | 9.97e-01 |
| **Overlap** | 9 |
| **Offset** | -14 |
| **Orientation** | Reverse Complement |
| | Show logo download options |

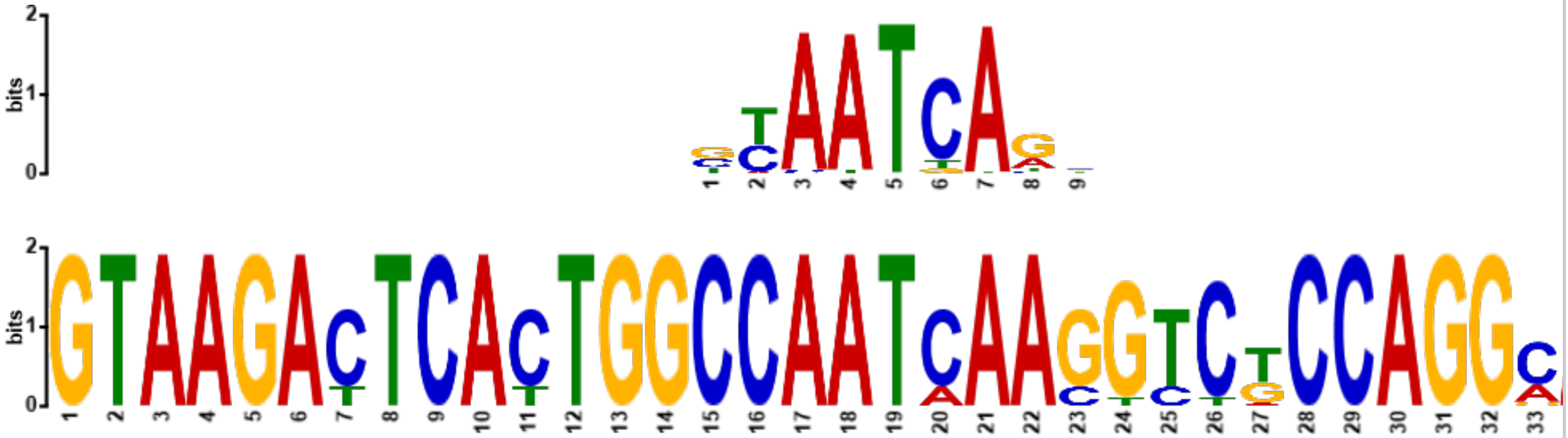

#### Settings

Previous Next Top

##### Alphabet

**Source:** the query file

| Name | Bg. | | | | Bg. | Name |
|---|---|---|---|---|---|---|
| Adenine | 0.245 | A | ~ | T | 0.245 | Thymine |
| Cytosine | 0.255 | C | ~ | G | 0.255 | Guanine |

##### Other Settings

| | |
|---|---|
| **Strand Handling** | Motifs may be reverse complemented before comparison to find a better match. |
| **Distance Measure** | Pearson correlation coefficient |
| **Match Threshold** | Matches must have a *E*-value of 10 or smaller. |

Previous Top

**Tomtom version**
5.4.1 (Release date: Sat Aug 21 19:23:23 2021 -0700)

**Command line**
```
tomtom -no-ssc -oc . -verbosity 1 -min-overlap 5 -mi 1 -dist pearson -evalue -thresh 10.0 -time 300 query_motifs db/MOUSE/HOCOMOCOv11_core_MOUSE_mono_meme_format.meme
```

Result calculation took 12.145 seconds
