## Supplemental_Figures_20231229 for "Imputation of 3D genome structure by genetic-epigenetic interaction modeling in mice"

### SUPPLEMENTAL INFORMATION

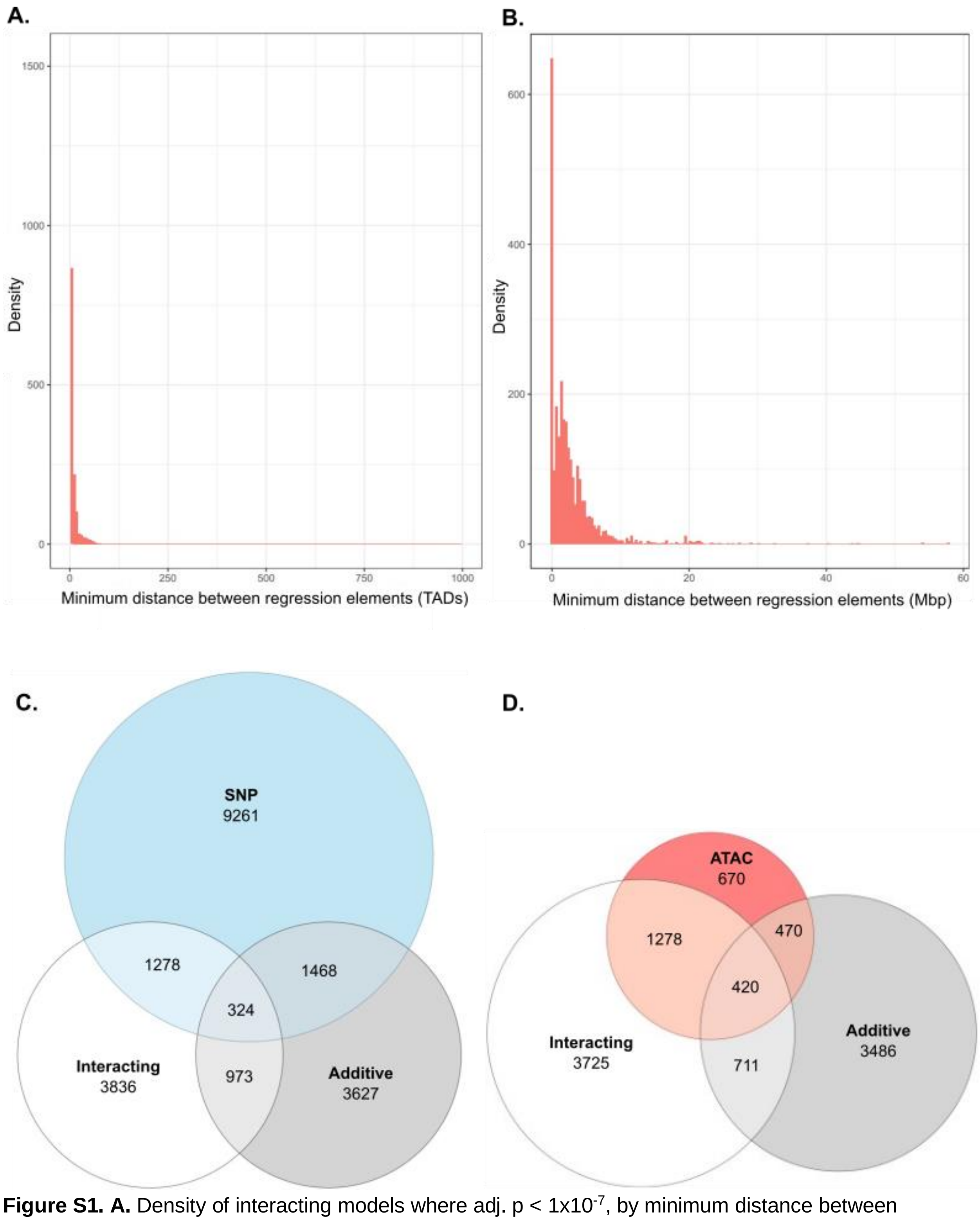


**Figure S1. A.** Density of interacting models where adj. $p < 1x10^{-7}$, by minimum distance between regulatory elements, measured in relation to number of TADs between elements. **B.** Density of additive models where adj. $p < 1x10^{-7}$, by minimum bp distance between regulatory elements. **C-D.** ATAC-seq peak and SNP participation in single-term, additive, and interacting models.

| Randomly generated sample database | Only SNP effect | Only ATAC effect | Additive effect | Interactive effect | No effect | All Models |
|---|---|---|---|---|---|---|
| **SNPs (p < 1x10^-7)** | 9261 | N/A | 3627 | 3836 | N/A | 68413 |
| **ATAC-seq peaks (p < 1x10^-7)** | N/A | 670 | 3486 | 3735 | N/A | 102104 |
| **Genes (p < 1x10^-7)** | 927 | 221 | 823 | 896 | 0 | 13631 |
| **Models** | 8059819 | 6952343 | 228616 | 560484 | 21166363 | 39021625 |
| **Models where p < 1x10^-7** | 14683 | 2465 | 4921 | 5110 | 0 | 27509 |
| **% Models where p < 1x10^-7** | 0.18% | 0.04% | 2.15% | 0.91% | 0.00% | 0.07% |

**Table S1.** Counts and percentages within a database of randomly generated regression models.

| TAD-focused sample database | Only SNP effect | Only ATAC effect | Additive effect | Interactive effect | No effect | All Models | All Models (no p cutoff) |
|---|---|---|---|---|---|---|---|
| **SNPs (p < 1x10^-7)** | 7247 | N/A | 12327 | 11904 | N/A | 15248 | 66193 |
| **ATAC-seq peaks (p < 1x10^-7)** | N/A | 487 | 22432 | 25961 | N/A | 53931 | 100751 |
| **Genes (p < 1x10^-7)** | 755 | 172 | 1129 | 1099 | N/A | 1223 | 13631 |
| **Models** | 47574072 | 20059818 | 16954276 | 14217610 | 52294007 | | 151123942 |
| **Models where p < 1x10^-7** | 750793 | 29117 | 305288 | 466451 | 0 | | 1551649 |
| **% Models where p < 1x10^-7** | 1.58% | 0.15% | 1.80% | 3.28% | 0.00% | | 1.03% |

**Table S2.** Counts and percentages within a database of all possible regression models where all SNPs and ATAC peaks are within +/- one TAD of the gene they interact with.

| | gene ~ geno | | gene ~ ATAC | | gene ~ geno + ATAC | | | | | gene ~ geno + ATAC + geno:ATAC | | | | |
|---|---|---|---|---|---|---|---|---|---|---|---|---|---|---|
| Chr | geno within +/- 2 Mb | % of total | ATAC within +/- 2 Mb | % of total | geno within +/- 2 Mb | % of total | ATAC within +/- 2 Mb | % of total | % proximal to each other? | geno within +/- 2 Mb | % of total | ATAC within +/- 2 Mb | % of total | % proximal to each other? |
| chr_1 | 709 | 54.16 | 4 | 80 | 117 | 54.17 | 6 | 2.78 | 3.24 | 126 | 45.49 | 15 | 5.42 | 10.11 |
| chr_10 | 345 | 43.56 | 0 | 0 | 54 | 39.42 | 9 | 6.57 | 3.65 | 53 | 50.48 | 4 | 3.81 | 3.81 |
| chr_11 | 508 | 62.95 | 0 | 0 | 75 | 63.56 | 10 | 8.47 | 0.85 | 101 | 54.01 | 10 | 5.35 | 8.02 |
| chr_12 | 203 | 64.24 | 1 | 100 | 34 | 59.65 | 1 | 1.75 | 0 | 15 | 41.67 | 5 | 13.89 | 13.89 |
| chr_13 | 246 | 57.34 | 3 | 100 | 48 | 55.17 | 7 | 8.05 | 4.60 | 52 | 36.88 | 10 | 7.09 | 14.18 |
| chr_14 | 282 | 43.79 | 56 | 71.79 | 55 | 43.65 | 23 | 18.25 | 6.35 | 39 | 22.94 | 45 | 26.47 | 8.24 |
| chr_15 | 207 | 65.30 | 0 | 0 | 51 | 72.86 | 4 | 5.71 | 2.86 | 26 | 50.98 | 6 | 11.76 | 7.84 |
| chr_16 | 218 | 53.56 | 0 | 0.00 | 40 | 44.94 | 4 | 4.49 | 3.37 | 27 | 39.71 | 3 | 4.41 | 10.29 |
| chr_17 | 400 | 55.94 | 4 | 50 | 95 | 53.37 | 14 | 7.87 | 4.49 | 62 | 47.33 | 17 | 12.98 | 7.63 |
| chr_18 | 97 | 43.50 | 0 | 0 | 27 | 52.94 | 4 | 7.84 | 5.88 | 18 | 29.03 | 4 | 6.45 | 8.06 |

| | | | | | | | | | | | | | | |
|---|---|---|---|---|---|---|---|---|---|---|---|---|---|---|
| chr_19 | 98 | 58.68 | 2 | 100 | 21 | 63.64 | 5 | 15.15 | 15.15 | 24 | 63.16 | 4 | 10.53 | 10.53 |
| chr_2 | 1251 | 48.77 | 75 | 75.76 | 175 | 44.08 | 46 | 11.59 | 6.80 | 209 | 39.36 | 79 | 14.88 | 8.10 |
| chr_3 | 449 | 46.00 | 2 | 50 | 111 | 45.87 | 7 | 2.89 | 4.96 | 75 | 38.46 | 7 | 3.59 | 14.87 |
| chr_4 | 642 | 57.22 | 4 | 100 | 162 | 67.78 | 13 | 5.44 | 1.67 | 95 | 51.63 | 8 | 4.35 | 4.89 |
| chr_5 | 709 | 53.63 | 2 | 66.67 | 139 | 55.16 | 8 | 3.17 | 2.78 | 92 | 40.00 | 10 | 4.35 | 5.22 |
| chr_6 | 266 | 47.58 | 0 | 0 | 32 | 49.23 | 3 | 4.62 | 6.15 | 38 | 33.63 | 7 | 6.19 | 10.62 |
| chr_7 | 493 | 53.41 | 4 | 80 | 108 | 59.67 | 6 | 3.31 | 3.31 | 77 | 48.43 | 4 | 2.52 | 5.03 |
| chr_8 | 221 | 58.47 | 15 | 68.18 | 38 | 46.91 | 7 | 8.64 | 2.47 | 51 | 43.97 | 19 | 16.38 | 6.03 |
| chr_9 | 278 | 58.77 | 3 | 100 | 52 | 57.78 | 5 | 5.56 | 4.44 | 35 | 44.87 | 7 | 8.97 | 14.10 |
| chr_X | 0 | 0 | 0 | 0 | 0 | 0 | 0 | 0 | 0 | 0 | 0 | 0 | 0 | 0 |
| **Totals** | 7622 | 51.34 | 175 | 52.12 | 1434 | 51.49 | 182 | 6.61 | 4.15% | 1215 | 41.10 | 264 | 8.47 | 8.57 |

**Table S3.** Counts and percentages of genotypic variants and ATAC-seq peaks within +/- 2 Mb of the gene they are imputed to affect. In additive and interacting models, we include the percent of models in which the genotypic variant and ATAC peak are closer to each other than to the gene they affect.

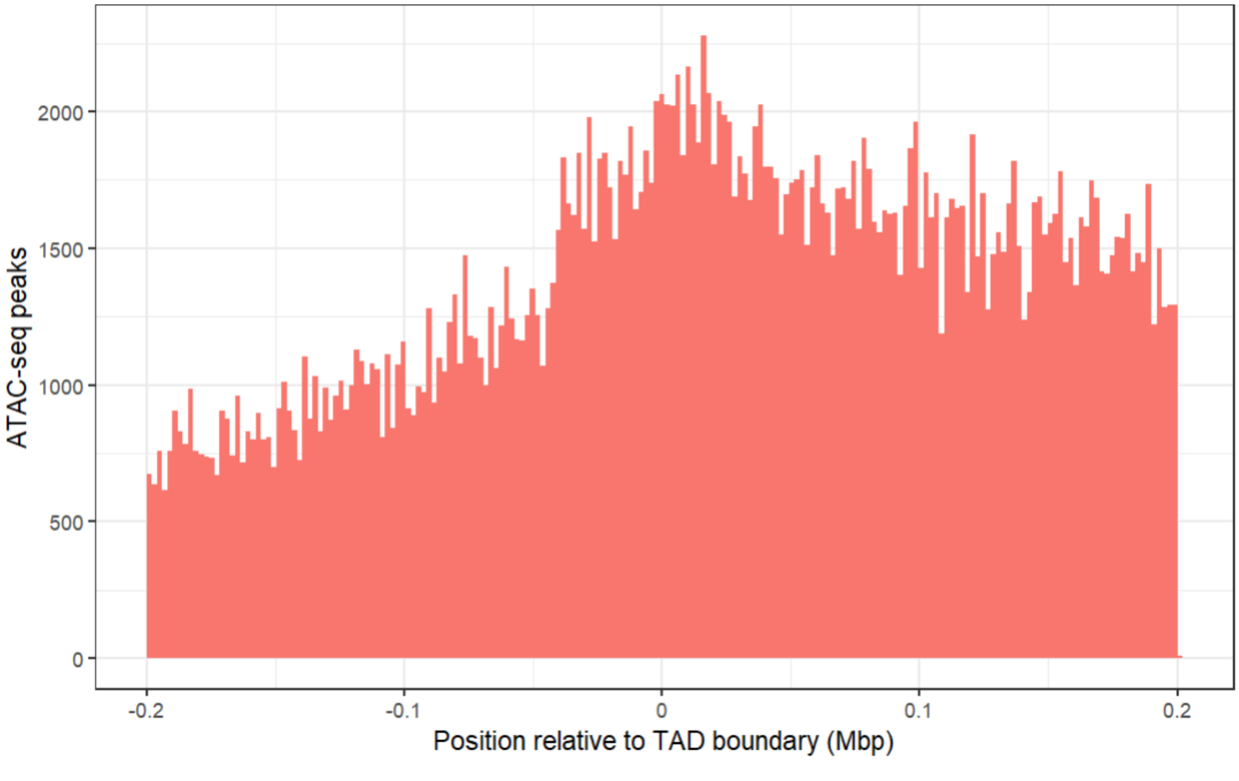


**Figure S2.** Location of all ATAC-seq peaks relative to TAD boundary location, merged across all genes.

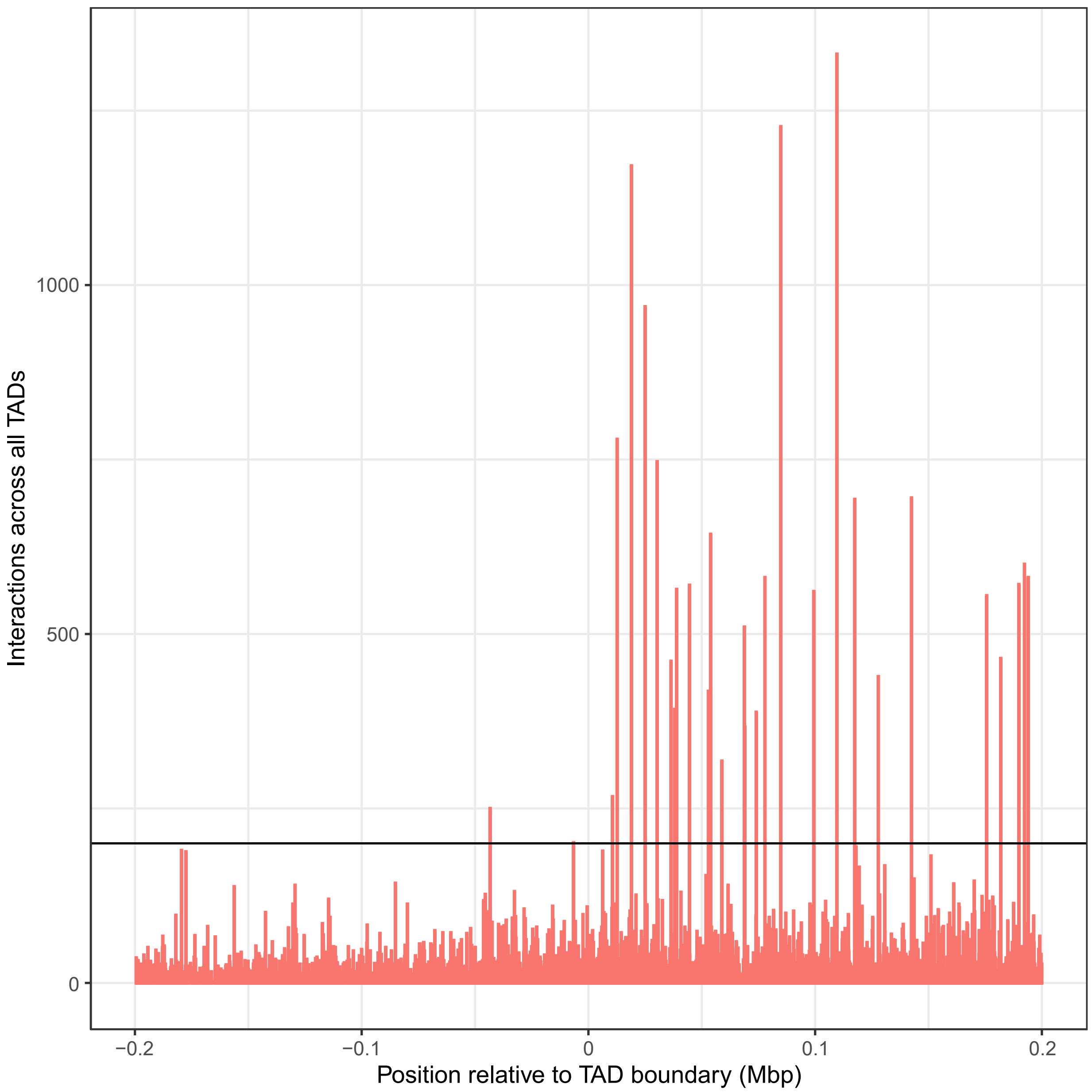

Interactions across all TADs
1000
500
0
−0.2
−0.1
0
0.1
0.2
Position relative to TAD boundary (Mbp)

**Figure S3.** Location of interacting ATAC-seq peaks relative to TAD boundary location, merged across all genes. Each bar represents an individual ATAC-seq peak, to demonstrate interactions per peak rather than density of interactions (see Fig 2B for this alternate view). Black horizontal line indicates 200 interactions per ATAC peak.

**A.**

| Chr | Count | In Transcript | % | In Intron | % | Before Coding | % | In Exon | % | After Coding | % |
|---|---|---|---|---|---|---|---|---|---|---|---|
| chr_1 | 1898 | 1231 | 64.86% | 1161 | 61.17% | 163 | 8.59% | 553 | 29.14% | 350 | 18.44% |
| chr_10 | 1210 | 842 | 69.59% | 803 | 66.36% | 157 | 12.98% | 460 | 38.02% | 240 | 19.83% |
| chr_11 | 1770 | 1263 | 71.36% | 1170 | 66.10% | 285 | 16.10% | 819 | 46.27% | 389 | 21.98% |
| chr_12 | 721 | 491 | 68.10% | 465 | 64.49% | 87 | 12.07% | 244 | 33.84% | 119 | 16.50% |
| chr_13 | 1101 | 626 | 56.86% | 594 | 53.95% | 88 | 7.99% | 281 | 25.52% | 166 | 15.08% |
| chr_14 | 1149 | 727 | 63.27% | 681 | 59.27% | 102 | 8.88% | 340 | 29.59% | 186 | 16.19% |
| chr_15 | 795 | 536 | 67.42% | 499 | 62.77% | 85 | 10.69% | 288 | 36.23% | 164 | 20.63% |
| chr_16 | 584 | 346 | 59.25% | 329 | 56.34% | 57 | 9.76% | 180 | 30.82% | 102 | 17.47% |
| chr_17 | 1852 | 1191 | 64.31% | 1139 | 61.50% | 196 | 10.58% | 592 | 31.97% | 301 | 16.25% |
| chr_18 | 627 | 425 | 67.78% | 412 | 65.71% | 59 | 9.41% | 207 | 33.01% | 74 | 11.80% |
| chr_19 | 660 | 476 | 72.12% | 459 | 69.55% | 103 | 15.61% | 263 | 39.85% | 165 | 25.00% |
| chr_2 | 2604 | 1706 | 65.51% | 1628 | 62.52% | 250 | 9.60% | 778 | 29.88% | 461 | 17.70% |
| chr_3 | 1425 | 915 | 64.21% | 850 | 59.65% | 173 | 12.14% | 494 | 34.67% | 287 | 20.14% |
| chr_4 | 1567 | 965 | 61.58% | 900 | 57.43% | 130 | 8.30% | 462 | 29.48% | 226 | 14.42% |
| chr_5 | 1886 | 1249 | 66.22% | 1184 | 62.78% | 202 | 10.71% | 618 | 32.77% | 338 | 17.92% |
| chr_6 | 1112 | 691 | 62.14% | 649 | 58.36% | 107 | 9.62% | 339 | 30.49% | 209 | 18.79% |
| chr_7 | 2039 | 1390 | 68.17% | 1303 | 63.90% | 274 | 13.44% | 864 | 42.37% | 454 | 22.27% |
| chr_8 | 1524 | 916 | 60.10% | 868 | 56.96% | 148 | 9.71% | 464 | 30.45% | 264 | 17.32% |
| chr_9 | 836 | 569 | 68.06% | 530 | 63.40% | 116 | 13.88% | 309 | 36.96% | 170 | 20.33% |
| chr_X | 601 | 399 | 66.39% | 372 | 61.90% | 97 | 16.14% | 282 | 46.92% | 160 | 26.62% |
| Total | 25961 | | | | | | | | | | |

| Chr | Enhancer | % | Promoter | % | Exon 1 | % | Single Exon | % | Outside | % |
|---|---|---|---|---|---|---|---|---|---|---|
| chr_1 | 113 | 5.95% | 362 | 19.07% | 233 | 12.28% | 14 | 0.74% | 609 | 32.09% |
| chr_10 | 69 | 5.70% | 339 | 28.02% | 207 | 17.11% | 11 | 0.91% | 334 | 27.60% |
| chr_11 | 106 | 5.99% | 538 | 30.40% | 369 | 20.85% | 28 | 1.58% | 456 | 25.76% |
| chr_12 | 27 | 3.74% | 170 | 23.58% | 120 | 16.64% | 8 | 1.11% | 217 | 30.10% |
| chr_13 | 43 | 3.91% | 205 | 18.62% | 121 | 10.99% | 11 | 1.00% | 442 | 40.15% |
| chr_14 | 34 | 2.96% | 222 | 19.32% | 151 | 13.14% | 10 | 0.87% | 388 | 33.77% |
| chr_15 | 49 | 6.16% | 186 | 23.40% | 126 | 15.85% | 9 | 1.13% | 232 | 29.18% |
| chr_16 | 124 | 21.23% | 126 | 21.58% | 73 | 12.50% | 9 | 1.54% | 193 | 33.05% |

| | | | | | | | | | | |
|---|---|---|---|---|---|---|---|---|---|---|
| chr_17 | 151 | 8.15% | 410 | 22.14% | 258 | 13.93% | 13 | 0.70% | 576 | 31.10% |
| chr_18 | 30 | 4.78% | 127 | 20.26% | 106 | 16.91% | 30 | 4.78% | 184 | 29.35% |
| chr_19 | 70 | 10.61% | 197 | 29.85% | 116 | 17.58% | 11 | 1.67% | 160 | 24.24% |
| chr_2 | 111 | 4.26% | 545 | 20.93% | 342 | 13.13% | 24 | 0.92% | 838 | 32.18% |
| chr_3 | 71 | 4.98% | 351 | 24.63% | 242 | 16.98% | 26 | 1.82% | 461 | 32.35% |
| chr_4 | 126 | 8.04% | 308 | 19.66% | 186 | 11.87% | 22 | 1.40% | 516 | 32.93% |
| chr_5 | 63 | 3.34% | 418 | 22.16% | 262 | 13.89% | 12 | 0.64% | 586 | 31.07% |
| chr_6 | 44 | 3.96% | 275 | 24.73% | 154 | 13.85% | 11 | 0.99% | 359 | 32.28% |
| chr_7 | 60 | 2.94% | 589 | 28.89% | 382 | 18.73% | 25 | 1.23% | 604 | 29.62% |
| chr_8 | 134 | 8.79% | 297 | 19.49% | 188 | 12.34% | 15 | 0.98% | 527 | 34.58% |
| chr_9 | 36 | 4.31% | 224 | 26.79% | 146 | 17.46% | 9 | 1.08% | 241 | 28.83% |
| chr_X | 12 | 2.00% | 227 | 37.77% | 131 | 21.80% | 15 | 2.50% | 182 | 30.28% |

**B.**

| | In Exons | In Exon 1 | In Enhancers | In Promoters | Anywhere Else |
|---|---|---|---|---|---|
| Count | 8837 | 3913 | 1473 | 6116 | 8105 |
| Percentage | 34.04% | 15.07% | 5.67% | 23.56% | 31.22% |
| Density per bp | 1.384 | 0.201 | 0.053 | 14.75 | 0.000 |

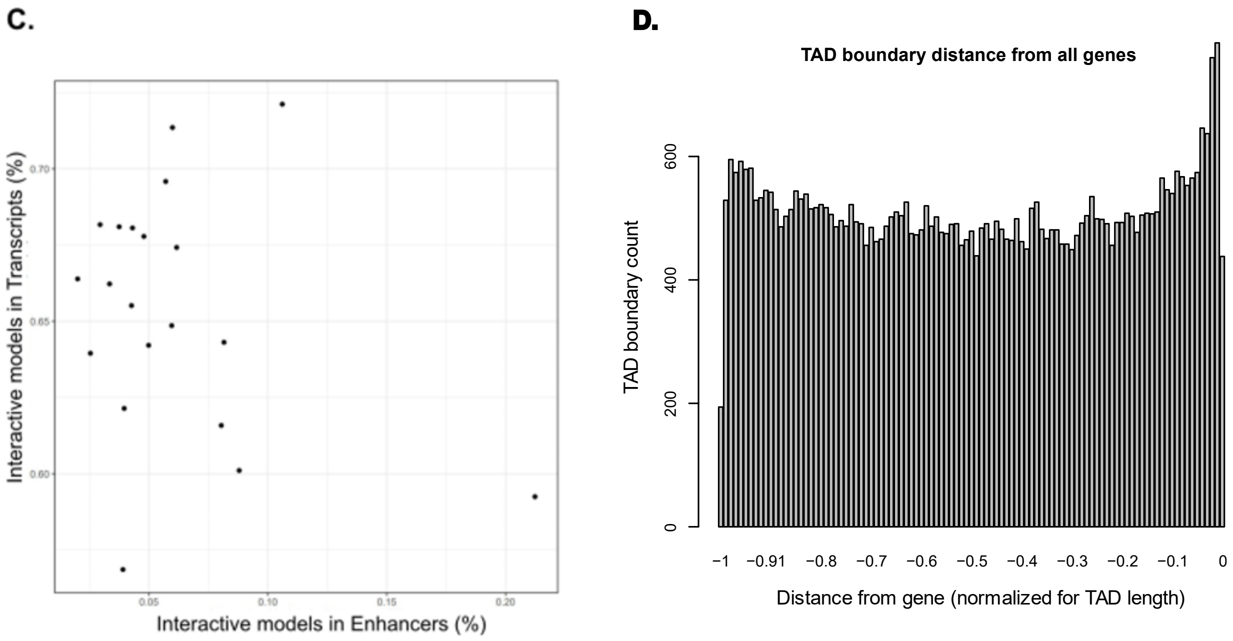


**Figure S4. A-B.** Tables providing a breakdown of interacting ATAC-seq peak locations relative to gene features **C.** Per-chromosome comparison of percentages of interacting ATAC-seq peaks in transcripts vs. in enhancers. **D.** TAD boundary locations relative to distance from each gene contained within them, normalized for TAD length.

**A.**

| Effect | Effect sign (all interacting models, adj. p < 1x10^-7) | | | | | | | |
|---|---|---|---|---|---|---|---|---|
| ATAC-seq | + | + | + | + | - | - | - | - |
| SNP | + | + | - | - | + | + | - | - |
| Interaction | + | - | + | - | + | - | + | - |
| % of total | 16.44% | 27.76% | 10.75% | 4.33% | 12.39% | 16.26% | 11.78% | 0.30% |

**B.**

| Effect | Effect sign (Platr2 interacting models, adj. p < 1x10^-7) | | | | | | | |
|---|---|---|---|---|---|---|---|---|
| ATAC-seq | + | + | + | + | - | - | - | - |
| SNP | + | + | - | - | + | + | - | - |
| Interaction | + | - | + | - | + | - | + | - |
| % of total | 4.31% | 38.55% | 1.13% | 12.47% | 12.47% | 9.07% | 21.77% | 0.23% |

**C.**

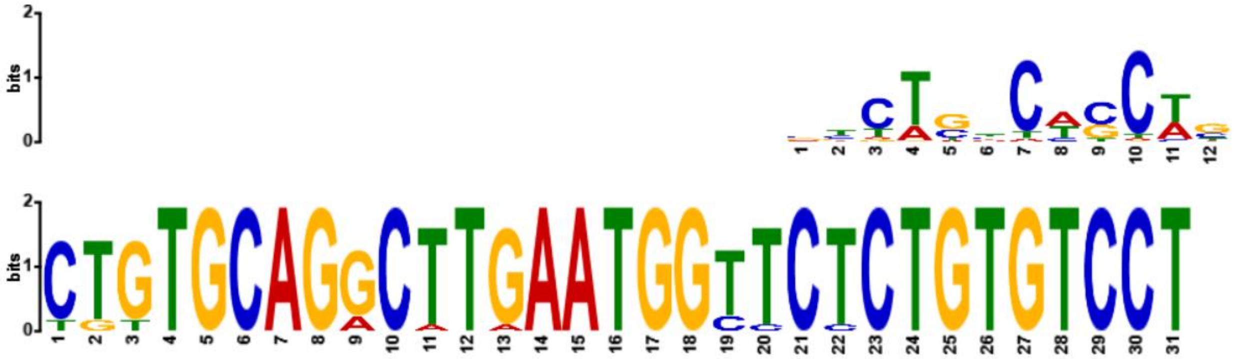

**D.**

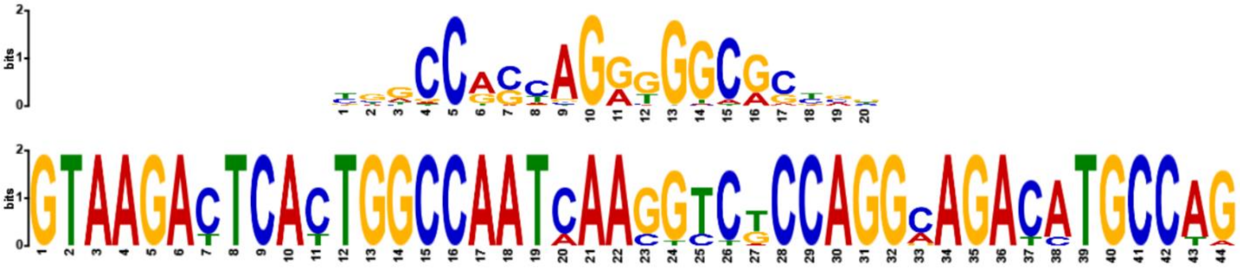

**Figure S5**. **A.** Model percentages calculated by distribution of effect signs for all significant interacting models. **B.** Platr2 interacting models **C-D.** Smad3 and Ctcf binding sites within motifs identified in Platr2's TAD

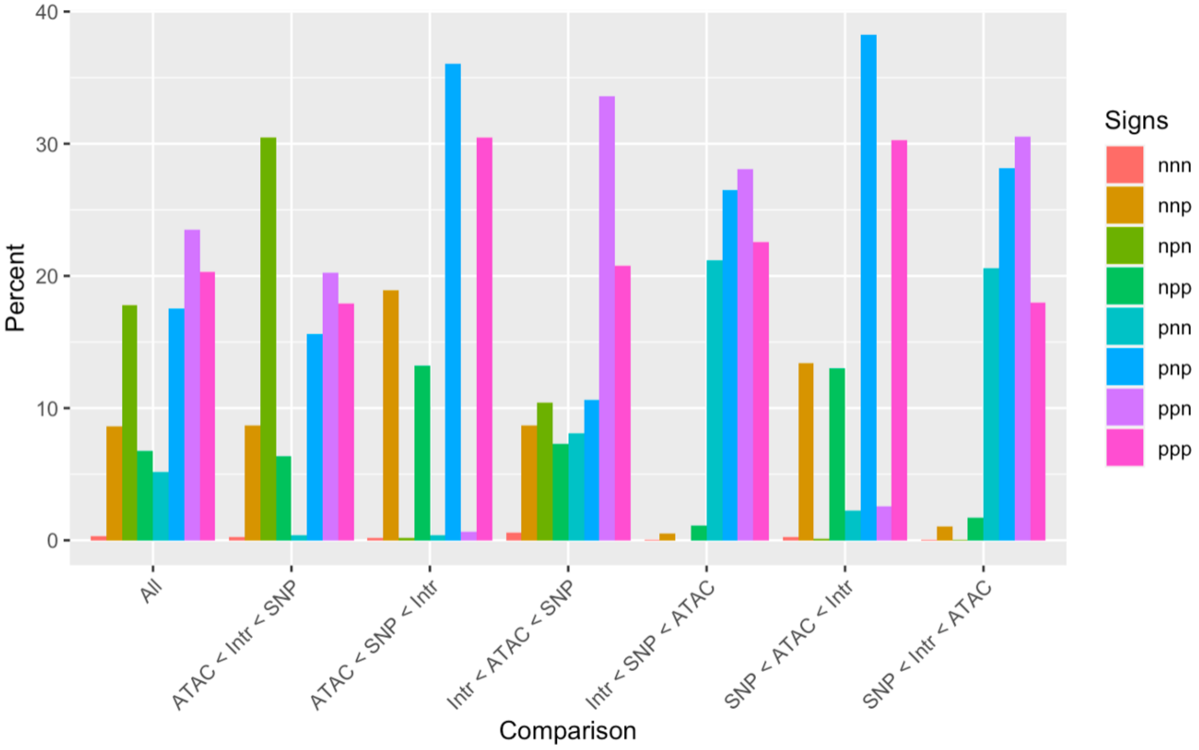


**Figure S6.** Relative effect magnitudes of all significant intra-TAD models, split by effect signs. Model signs are listed in the order ATAC, SNP, Interaction, and positive (p) or negative (n).

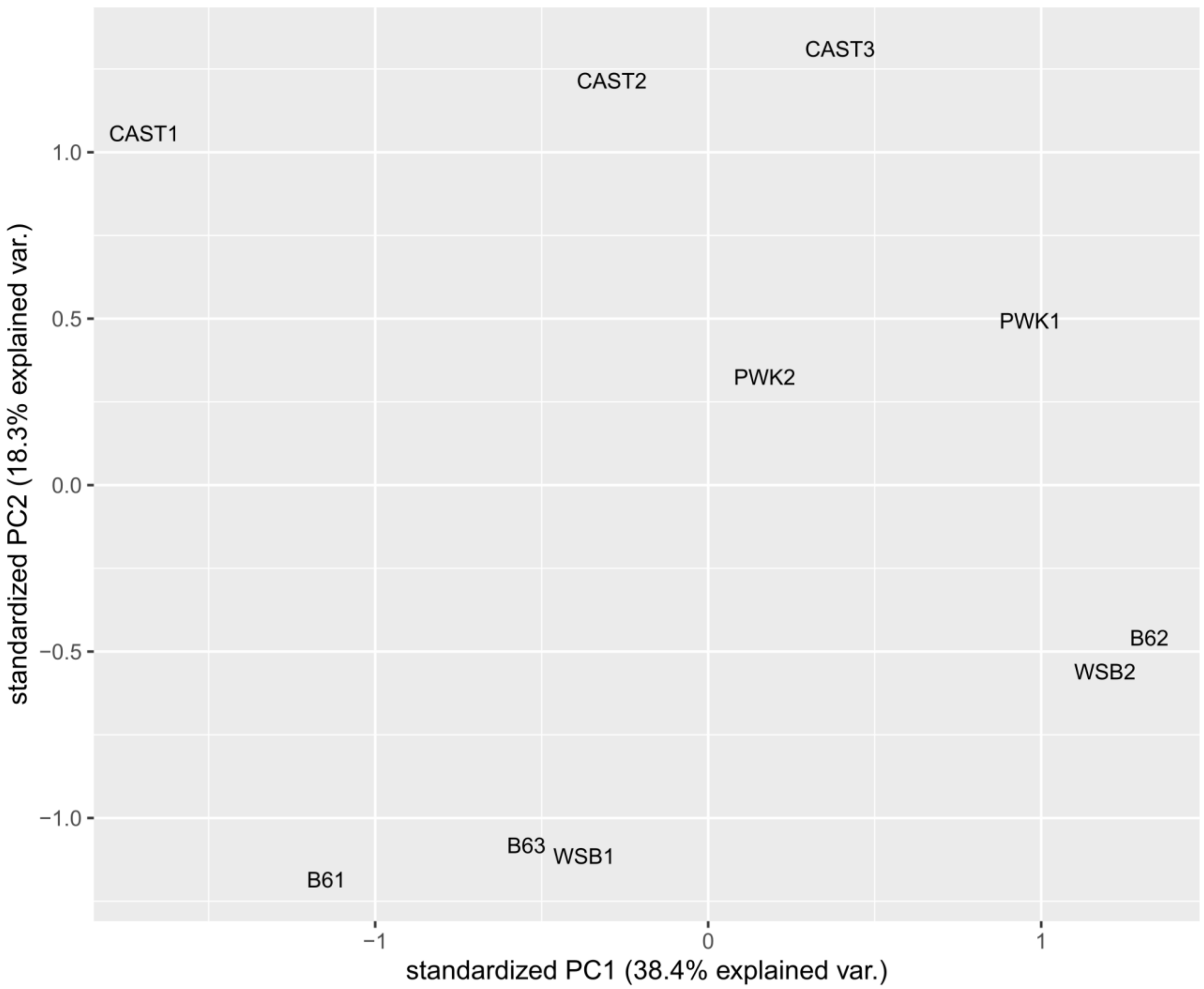


**Figure S7.** Principle component analysis of CTCF ChIP-seq binding intensity.
